# Energetic costs, precision, and efficiency of a biological motor in cargo transport

**DOI:** 10.1101/200907

**Authors:** Wonseok Hwang, Changbong Hyeon

## Abstract

Molecular motors play key roles in organizing the interior of cells. An efficient motor in cargo transport would travel with a high speed and a minimal error in transport time (or distance) while consuming minimal amount of energy. The travel distance and its variance of motor are, however, physically constrained by energy consumption, the principle of which has recently been formulated into the *thermodynamic uncertainty relation*. Here, we reinterpret the uncertainty measure (𝒬) defined in the thermodynamic uncertainty relation such that a motor efficient in cargo transport is characterized with a small 𝒬. Analyses on the motility data from several types of molecular motors show that 𝒬 is a nonmonotic function of ATP concentration and load (*f*). For kinesin-1, 𝒬 is locally minimized at [ATP] ≈ 200 *μ*M and *f* ≈ 4 pN. Remarkably, for the mutant with a longer neck-linker this local minimum vanishes, and the energetic cost to achieve the same precision as the wild-type increases significantly, which underscores the importance of molecular structure in transport properties. For the biological motors studied here, their value of 𝒬 is semi-optimized under the cellular condition ([ATP] ≈ 1 mM, *f* = 0 − 1 pN). We find that among the motors, kinesin-1 at single molecule level is the most efficient in cargo transport.

---

Biological systems are in nonequilibrium steady states (NESS) in which the energy and material currents flow constantly in and out of the system. Subjected to incessant thermal and nonequilibrium fluctuations, cellular processes are inherently stochastic and error-prone. To minimize detrimentaion effects, cells are equipped with a plethora of energy-consuming error-correction mechanisms. Trade-off relations between the energetic cost and information processing are ubiquitous in cellular processes, and have been a recurring theme in biology for many decades [1? ? –4].

A recent study by Barato and Seifert [5] has formulated a concise inequality known as the *thermodynamic uncertainty relation*, the trade-off between energy consumption and precision of an observable from dissipative processes in NESS. To be more explicit, the uncertainty measure 𝒬, defined as a product between the energy consumption (heat dissipation, *Q*(*t*)) from an energy-driven process in the steady state and the squared relative error of an output observable *X*(*t*), 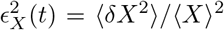, is constant. It has been further conjectured that for an arbitrary chemical network formulated by Markov jump processes 𝒬 cannot be smaller than 2*k*_*B*_*T*,

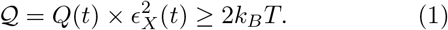

The measure 𝒬 quantifies the uncertainty of a dynamic process. The smaller the value of 𝒬, the more regular and predictable is the trajectory generated from the process, rendering the output observable more precise. The proof and physical significance of this inequality have been discussed [5–10]. In the presence of large fluctuations inherent to cellular processes, harnessing energy into precise motion is critical for accuracy in cellular computation. The uncertainty measure 𝒬 can be used to assess the efficiency of suppressing the uncertainty in dynamical process via energy consumption.

Historically, the efficiency of heat engine has been discussed in terms of the thermodynamic efficiency, the aim of which is to maximize the amount of work extracted from heat reservoirs with two different temperatures [11]. For a nonequilibrium machine powered by chemical driving forces constantly regulated in the live cell, power production for a given free energy consumption could be a relevant quantity to maximize. Meanwhile, for transport motors in cells, the uncertainty measure 𝒬 can be used to assess the efficiency of a motor (or motors) in cargo transport. Because the displacement (or travel distance) is a natural output observable of interest in the cargo transport, we set *X* = *l*(*t*), which recasts Eq.1 into

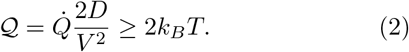

𝒬 is minimized by a motor that transports cargos (i) at high speed (*V* ~ 〈*l*(*t*)〉/*t*), (ii) with small error (*D* ~ 〈*δl*(*t*)^2^〉/*t*) in the displacement (or punctual delivery to a target site), and (iii) with small energy consumption 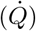. An efficient cargo transporter is characterized by a small 𝒬 with its minimal bound 2*k*_*B*_*T*.

Here, we assess the “efficiency” of several biological motors (kinesin-1, KIF17 and KIF3AB in kinesin-2 family, myosin-V, dynein, and F_1_-ATPase) in terms of 𝒬, and study how it changes with varying conditions of load (*f*) and [ATP]. Also, of great interest is to identify the optimal condition for motor efficiency, if any, that 𝒬 is minimized. To evaluate 𝒬, one should know 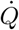, *D*, and *V* of the system (see Eq.2), which can be obtained usinga proper kinetic network model that can delineate the dynamical characteristics of the system [12]. Our analyses on motors show that 𝒬 is a complex function of f and [ATP] and is sensitive to a subtle variation in motor structure. Transport and rotory motors studied here are semi-optimized in terms of 𝒬 under cellular condition.

## RESULTS AND DISCUSSION

### Chemical driving force, steady state current, and heat dissipation of double-cycle kinetic network for kinesin-1

The chemomechanical dynamics of a molecular motor can be mapped on an appropriate kinetic network, which allows us to express the transport properties of the motor, *V* and *D*, in terms of a set of rate constants {*k*_*ij*_}, where *k*_*ij*_ is associated with the transition from the *i*-th to *j*-th state [5, 13, 14] (see **SI** text). The formal expressions of *V* and *D* in terms of {*k*_*ij*_(*f*, [ATP])} are used to fit simultaneously the experimental data of *V* and *D* obtained under varying conditions of ATP and *f* [12]. The set of rate constants {*k*_*ij*_} decided from the fit further allows us to calculate the affinity (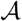, net driving force), and reaction current (*J*) of the network, and hence the heat dissipation from the motor, 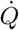 (see below).

For kinesin-1, we considered the 6-state network [15], consisting of two cycles, 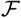 and 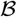 (Fig. 1A). This minimal kinetic scheme accommodates 4 different kinetic paths: ATP-hydrolysis induced forward/backward step and ATP-synthesis induced forward/backward step. Although the conventional (N=4)-state unicyclic kinetic model confers a similar result with the 6-state double-cycle network model at small *f* (compare Figs.1 and Fig.S3), it is led to a physically problematic interpretation when the molecular motor is stalled and starts taking backsteps at large hindering load. The backstep in unicyclic network, by construction, is produced by a reversal of the forward cycle [16], which implies that the backstep is always realized via the synthesis of ATP from ADP and P_*i*_. More importantly, in calculating 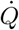 from kinetic network, the unicyclic network results in 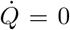 under the stall condition, which is not compatible with the physical reality that an idling car still burns fuel and dissipates heat 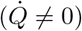. To build a more physically sensible model that considers the possibility of ATP-induced (fuel-burning) backstep, we extend the unicyclic network into a multi-cyclic one which takes into account an ATP-consuming stall, i.e., a futile cycle [15–18].

**FIG. 1.**
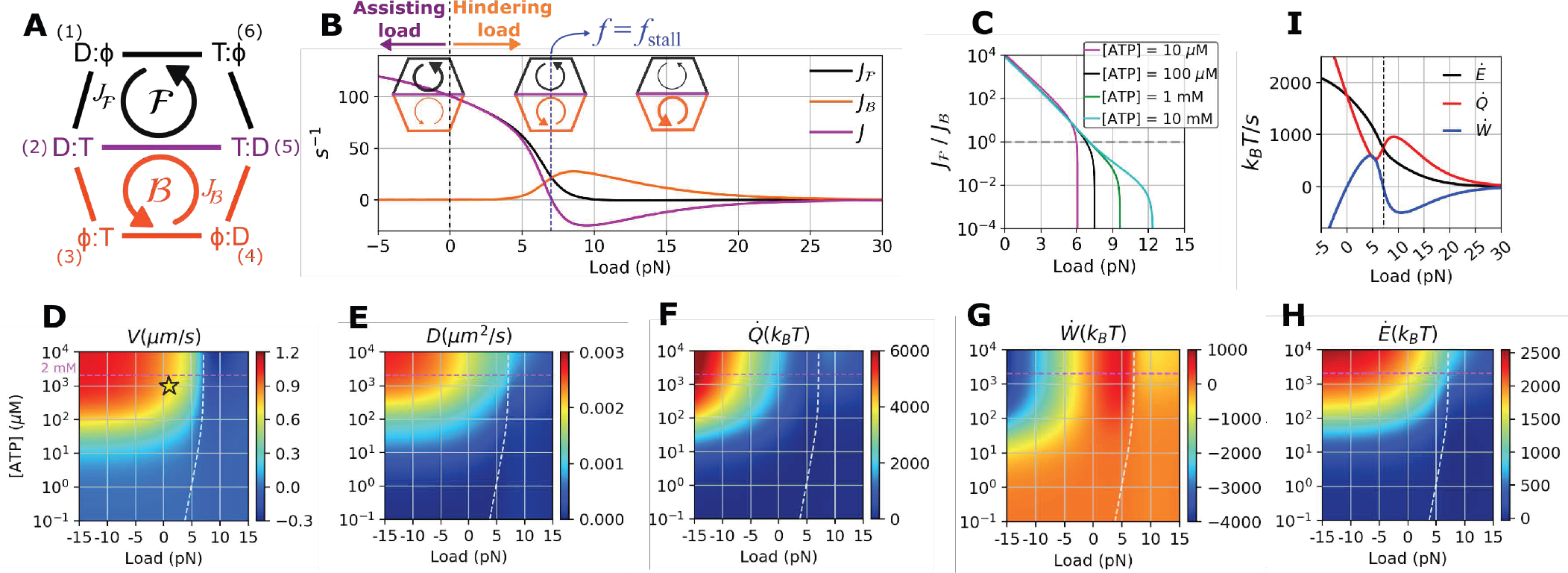
Analysis of kinesin-1 based on the 6-state double-cycle kinetic model. **A.** Schematics of the 6-state double-cycle kinetic network for hand-over-hand dynamics of kinesin-1, where *T*, *D*, and *ϕ* denote ATP-, ADP-bound, and apo state, respectively. Through binding [(1)→(2)] and hydrolysis of ATP [(2)→(5)], and release of ADP [(5)→(6)], kinesin moves forward in the 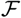-cycle [(1) → (2) → (5) → (6) → (1)], where it takes a backstep in the 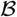-cycle [(4) → (5) → (2) → (3) → (4)]. The arrows in the figure depict the direction of reaction currents. In both cycles, each chemical step is reversible and the transition rate from the state (*i*) to (*j*) is given by *k*_*ij*_. **B.** Reaction current 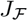, 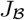, and *J* as a function of load. The three cartoons illustrate the amount of current along the 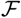 and 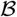 cycles at each value of load. *f* < 0 and *f* > 0 correspond to the assisting and hindering load, respectively. **C.** The ratio between the forward and backward uxes 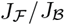 as function of *f* at fixed [ATP]. The stall forces, determined at 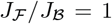 (dashed line), are narrowly distributed between *f* = 6 − 8 pN. (**D**−**H**) *V*, *D*, 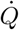, *Ẇ*, *Ė* as a function of f and [ATP]. The white dashed lines depict the locus of [ATP]-dependent stall force, and the dashed lines in magenta indicate the condition of [ATP] = 2 mM. **I.** Dependences of *Ė*, 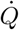, *Ẇ* on *f* at [ATP] = 2 mM. The star symbol in **D** indicates the cellular condition of *f* ≈ 1 pN and [ATP] ≈ 1 mM, which will be discussed later.

In the double-cycle model, kinesin-1 predominantly moves forward through 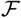-cycle under small hindering (*f* > 0) or assisting load (*f* < 0), whereas it takes a backstep through the 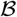-cycle under large hindering load. In principle, the current within the 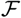-cycle, 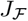, itself is decomposed into the forward 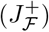 and backward current 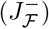, and 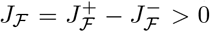 is satisfied in NESS. Although a backstep could be realized through an ATP synthesis [19], corresponding to 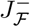, experimental data [16, 20, 21] suggest that such backstep current (ATP synthesis induced backstep, 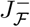) is negligible in comparison with 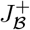 (ATP hydrolysis induced backstep).

The dynamics realized in the double-cycle network can be illuminated in terms of variation of 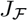 and 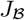 with increasing *f* (Fig.1B). Without load (*f* = 0), kinesin-1 predominantly moves forward 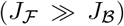. This imbalance diminishes with increasing *f*. At stall conditions, the two reaction currents are balanced 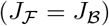, so that the net current *J* associated with the mechanical stepping defined between the states (2) and (5) vanishes 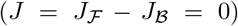, but nonvanishing current due to chemistry still remains along the cycle of → (2) → (3) → (4) → (5) → (6) → (1) → (2) →. A further increase of f beyond the stall force renders 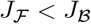, augmenting the likelihood of backstep.

For a given set of rate constants, it is straightforward to calculate the rates of heat dissipation 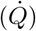, work production (*Ẇ*), and total energy supply (*Ė*). The total heat generated from the kinetic cycle depicted in Fig. 1A is decomposed into the heat generated from two subcycles, 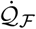 and 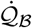, each of which is the product of reaction current and affinity [5, 15, 22–25]

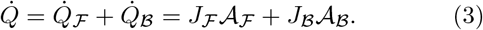

Here, the affinities (driving forces) for the 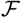 and 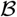 cycles are

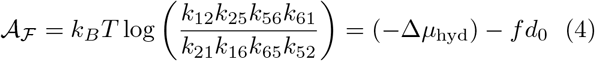

and

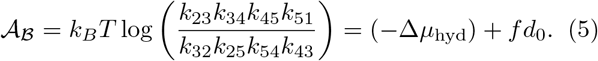

The explicit forms of 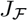 as a function of {*k*_*ij*_} are available (see Eq. S25) but the expression is generally more complicated than the affinity. It is of note that at *f* = 0, the chemical driving forces for 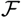 and 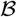 cycles are identical to be –Δ*μ*_hyd_. The above decomposition of affinity associated with each cycle into the chemical driving force and the work done by the motor results from the Bell-like expression of transition rate between the states (2) and (5) as 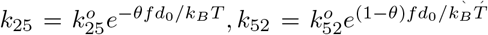 [15, 26, 27] (see **Methods**). Thus, Eqs. 3, 4, and 5 lead to

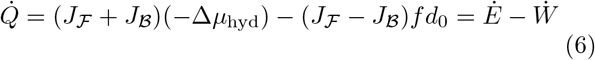

where 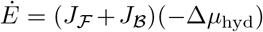 is the rate of total chemical free energy supplied to the system via ATP hydrolysis. 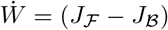*fd*_0_ is the rate of work production.

Dependences of *V*, *D*, 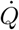, and *Ẇ* for kinesin-1 on *f* and [ATP] make a few points visually clear (see Fig. 1): (i) The stall condition, indicated with a white dashed line in each map, divided all the 2D maps of *V*, *D*, 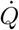, and *Ẇ* into two regions; (ii) In contrast to *V* (Fig. 1D) and *D* (Fig. 1E), which decrease monotonically with *f*, 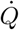 and *Ẇ* display non-monotonic dependence on *f* (Figs. 1F, G). At high [ATP], 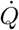 is maximized at *f* = 0, 10 pN, whereas *Ẇ* is maximized at 5 pN and minimized at 10 pN (see Fig. 1I calculated at [ATP]=2 mM).

At the stall force *f*_stall_ (≈ 7 pN) (Figs. 1B, 1I, black dashed line), the reaction current of the 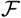-cycle is exactly balanced with that of 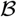-cycle 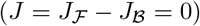, giving rise to zero work production (*Ẇ* = *fV* = *fd*_0_*J* = 0). The numbers of forward and backward steps taken by kinesin motors are identical, and hence there is no net directional movement (*V* = 0) [21]. It is important to note that even at the stall condition, kinesin-1 consumes the chemical free energy of ATP hydrolysis, dissipating heat in both forward and backward steps, and hence rendering 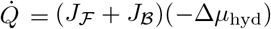 always positive.

Next, branching of reaction current of the 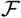-cycle into 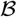-cycle with increasing f gives rise to nonmonotonic changes of 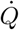 and *Ẇ* with *f*. At small *f*, 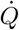 decreases with *f* because exertion of load gradually deactivates the 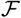-cycle via the decrease of 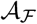 (Eq.3, Eq.4, Fig. S2C) and 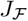 (Fig. S2A). By contrast, 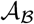 and 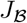 increase with *f* (Eq.5, Figs. S2A and D), which leads to an increase of 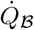. The nonmonotic dependence of *Ẇ* on *f* can be analyzed in a similar way. Because the heat dissipated from the whole network is decomposed into the parts 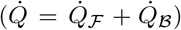, the activation and deactivation of the cycles 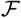 and 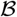 (i.e., 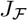 and 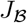) are negatively correlated (Figs. S2A and B).

**FIG. 2.**
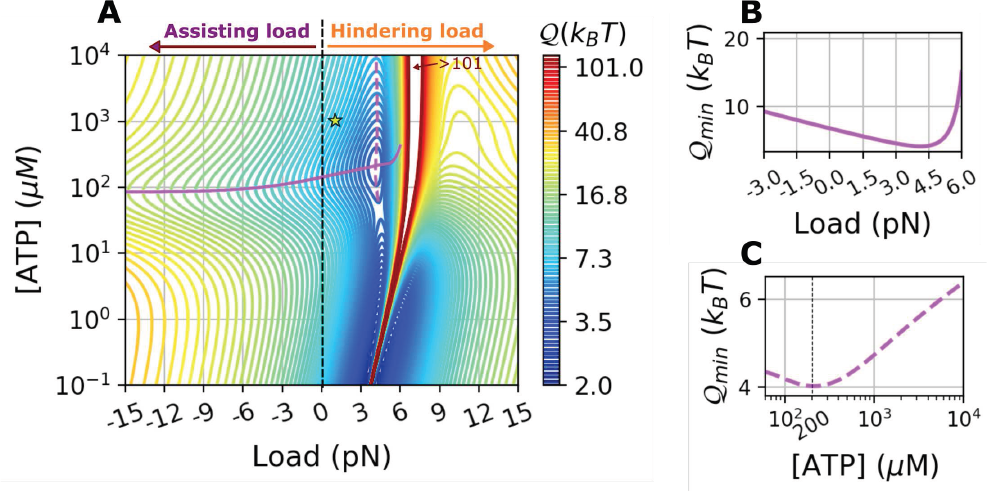
𝒬 calculated based on kinesin-1 data [31] using the 6-state double-cycle model [15] at varying *f* and [ATP], where *f* > 0 and *f* < 0 signify the hindering and assisting load, respectively. **A.** 2-D contour plot of 𝒬 = 𝒬(*f*, [ATP]). The solid lines in magenta are the loci of minimal 𝒬 at varying [ATP] for a given *f* (**B**). The dashed lines in magenta are the loci of minimal 𝒬 at varying *f* for a given [ATP] (**C**). The star symbol indicates the cellular condition of [ATP] ≈ 1 mM and *f* ≈ 1 pN.

Motor head distortion at high external stress hinders the binding and hydrolysis of ATP in the catalytic site [28–30]; *k*_*ij*_ = 0 when ATP cannot be processed. This effect is modeled into the rate constants such that 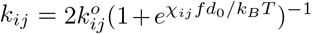 with *χ_ij_* > 0 (*k*_*ij*_ ≠ *k*_25_, *k*_52_) [15]. Thus, it naturally follows that 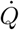 = 0 at f far greater than f_stall_ (Fig.1B, F, I).

### Quantification of 𝒬 for kinesin-1

Unlike *V*, *D*, and 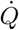 that are maximized at large [ATP] and small *f* (Figs. 1D, E, F), the uncertainty measure 𝒬(*f*, [ATP]) displays a complex functional dependence (Fig. 2A). (i) Small 𝒬 at low [ATP] is a trivial outcome of the detailed balance condition, when the amount of [ATP] is in balance with [ADP] and [P_*i*_]. The motor, without chemical driving force and only subjected to thermal fluctuations 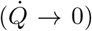, is on average motionless (*V* → 0); 𝒬 is minimized in this case (𝒬 → 2*k*_*B*_*T*. (ii) 𝒬 is generally smaller below the stall condition, *f* < *f*_stall_([ATP]), demarcated by the white dashed lines in Fig.1. In this case, the reaction current along the 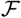-cycle is more dominant than that above the stall. At the stall, 𝒬 diverges because of *V* → 0 and 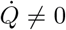. (iii) Notably, a suboptimal value of 𝒬 ≈ 4 *k*_B_*T* is identified at [ATP] = 210 *μ*M and *f* = 4.1 pN (Figs. 2).

### Comparison of 𝒬 between different types of kinesins

The dynamic property of molecular motor differs from one motor type to another. Effect of modifying motor structure on the transport properties as well as on the directionality and processivity of molecular motor has been of great interest because it provides glimpses into the design principle of a motor at molecular level [30, 33–36]. Here, we explore how modification to motor structure changes the efficiency of motor in terms of 𝒬. To this end, we analyze single-molecule motility data of a mutant of kinesin-1 (Kin6AA), and homodimeric and heterotrimeric kinesin-2 (KIF17 and KIF3AB).

The data of Kin6AA, a mutant of kinesin-1 that has longer neck-linker domain in each monomer, were taken from Ref. [18]. Six amino-acid residues inserted to the neck-linker reduces the internal tension along the neck-linker which plays a critical role for regulating the chemistry of two motor heads and coordinating the handover-hand motion [16, 37]. Disturbance to this motif is expected to affect *V* and *D* of the wild type. We analyzed the data of Kin6AA again using the 6-state network model (Figs. 1A, S4, S5, Table II. See **SI** for detail), indeed finding reduction of *V* and *D* (Fig. S5A) as well as its stall force (Fig. S5A, white dashed line). Of particular note is that the rate constant *k*_25_ associated with the mechanical stepping process is reduced by two order of magnitude (Table II). In 𝒬(*f*, [ATP]) (Fig. 3A), the suboptimal point observed in kinesin-1 (Fig. 2A) vanishes (Fig. 3A), and 𝒬 diverges around ~ 4 pN due to the decreased stall force. Finally, overall, the value of 𝒬 has increased dramatically. This means that compared with kinesin-1 (𝒬 ≈ 7 *k*_*B*_*T*, the trajectory of Kin6AA is less regular and unpredictable (𝒬 ≈ 20 *k*_*B*_*T*), and thus Kin6AA is three fold less efficient in cargo transport. Note that 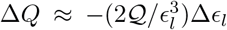, and hence it is energetically demanding to improve the precision when *∊*_*l*_ is already small.

**FIG. 3.**
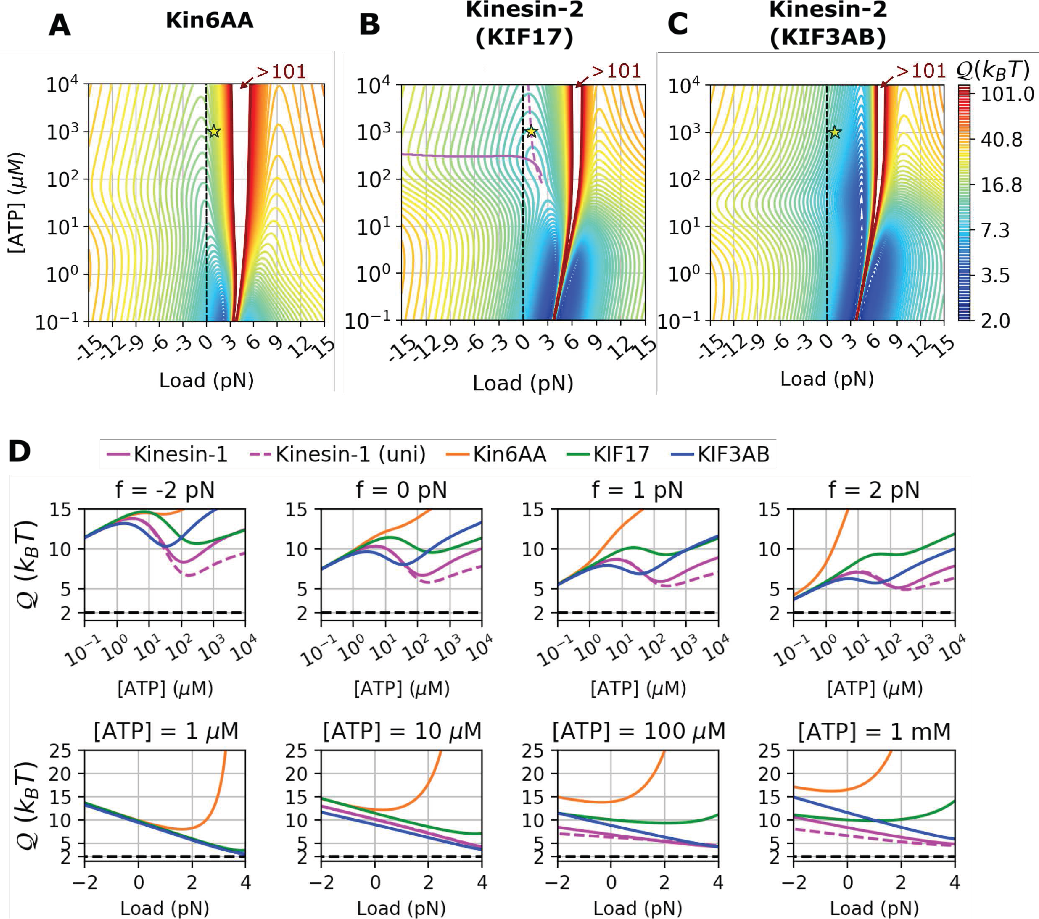
FIG. 3. 𝒬(*f*, [ATP]) calculated for (**A**) Kin6AA [18], (**B**) homodirmeric kinesin-2 KIF17 [32], and (**C**) heterotrimeric kinesin-2 KIF3AB [32]. In **A**-**C**, the condition of [ATP] ≈ 1 mM and *f* ≈ 1 pN is indicated with the star symbols. **D**. 𝒬([ATP]) at fixed *f* (upper panels) and 𝒬(*f*) at fixed [ATP] (lower panels) calculated for kinesin-1 (solid magenta lines for 6-state network model and dashed magenta lines for unicyclic model), Kin6AA (orange lines), KIF17 (green lines), and KIF3AB (blue lines). 𝒬 = 2*k*_*B*_*T* is marked with the black dashed lines.

Next, the values of 𝒬 were calculated for two active forms of vertebrate kinesin-2 class motors responsible for intraflagellar transport (IFT). KIF17 is a homodimeric form of kinesin-2, and KIF3AB is a heterotrimeric form made of KIF3A, KIF3B, and a nonmotor accessory protein, KAP. To quantify their motility properties, we digitized single-molecule motility data from Ref. [32] and fitted them to the 6-state double-cycle model (Figs. S6, S8) (See **SI** for detail). 𝒬(*f*, [ATP]) of KIF17 and KIF3AB are qualitatively similar to that of kinesin-1 with some variations. 𝒬 for KIF17 forms a shallow local minimum of 𝒬 ≈ 9.2 *k*_*B*_*T* at [ATP] = 200 *μ*M and *f* = 1.5 pN (Fig. 3B), whereas such suboptimal condition vanishes in KIF3AB (Fig. 3C). However, KIF3AB has a local valley around f ~ 4 pN and 1 *μ*M ≲ [ATP] ≲ 10 mM in which 𝒬 ≈ 4*k*_*B*_*T*.

The plots of 𝒬([ATP]) at fixed *f* and 𝒬(*f*) with fixed [ATP] in Fig. 3D recapitulate the difference between different classes of kinesins more clearly. The following features are noteworthy. (i) An extension of neck-linker domain (Kin6AA, orange lines) dramatically increases 𝒬 compared with the wild type (Kinesin WT, magenta lines). (ii) Non-monotonic behaviors of 𝒬 are qualitatively similar for all kinesins although 𝒬 is, in general, the smallest for kinesin-1. (iii) The movement of KIF3AB (black lines) becomes the most regular at low [ATP] (≲ 10 *μ*M). (iv) 𝒬 for the unicyclic model of kinesin-1 (dashed magenta lines) displays only small deviations as long as 0 ≲ *f* ≲ 4 pN ≪ *f*_stall_ ≈ 7 pN.

### Comparison of 𝒬 among different types of motors

We further investigate 𝒬(*f*, [ATP]) for myosin-V, dynein, and F_1_-ATPase using the kinetic network models proposed in the literature [38–40].

#### Myosin-V

The model studied in Ref. [38] consists of chemomechanical forward cycle 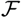, dissipative cycle *ε*, and pure mechanical cycle 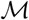 (Fig. 4A). In 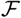-cycle, myosin-V either moves forward by hydrolyzing ATP or takes backstep via ATP synthesis. In 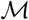-cycle, myosin-V moves backward under load without involving chemical reactions. The *ε*-cycle, consisting of ATP binding [(2) → (5)], ATP hydrolysis [(5) → (6)], and ADP release [(6) → (2)] (Fig. 4B), was originally introduced to connect the two cycles 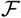 and 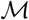. The calculation of 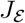 and 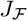 reveals that a gradual deactivation of 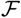-cycle with decreasing [ATP] activates the *ε*-cycle (Fig. S11A, 100 *μ*M ≲ [ATP] ≲ 1 mM, *f* ≲ 1 pN). Thus, *ε*-cycle can be regarded a futile 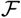-cycle, which is activated when chemical driving force is balanced with a load *f* at low [ATP]. 𝒬(*f*, [ATP]) calculated at [ADP] = 70 *μ*M and [P_*i*_] = 1 mM using the rate constants from Ref. [38] (see **SI** for details and Fig. S11) reveals no local minimum in this condition. However, at [ADP] =0.1 *μ*M and [P_*i*_] = 0.1 *μ*M, which is the condition studied in Ref. [38], a local minimum with 𝒬 = 6.5 *k*_*B*_*T* emerges at *f* = 1.1 pN and [ATP] = 20 *μ*M (Fig. S12D, Table I). Both values of *f* and [ATP] at the suboptimal condition of myosin-V are smaller than those of kinesin-1 (Table I). In (N=2)-unicyclic model for myosin-V (Fig. S15) [41], 𝒬 has local valley around *f* ~ 2 pN and [ATP] ~ 10 *μ*M. Similar to the result from the multi-cyclic model with [ADP] =0.1 *μ*M and [P_*i*_] =0.1 *μ*M, the values of *f* and [ATP] along the valley of 𝒬 are smaller than the values optimizing 𝒬 for the kinesin-1 (Table I).

**FIG. 4.**
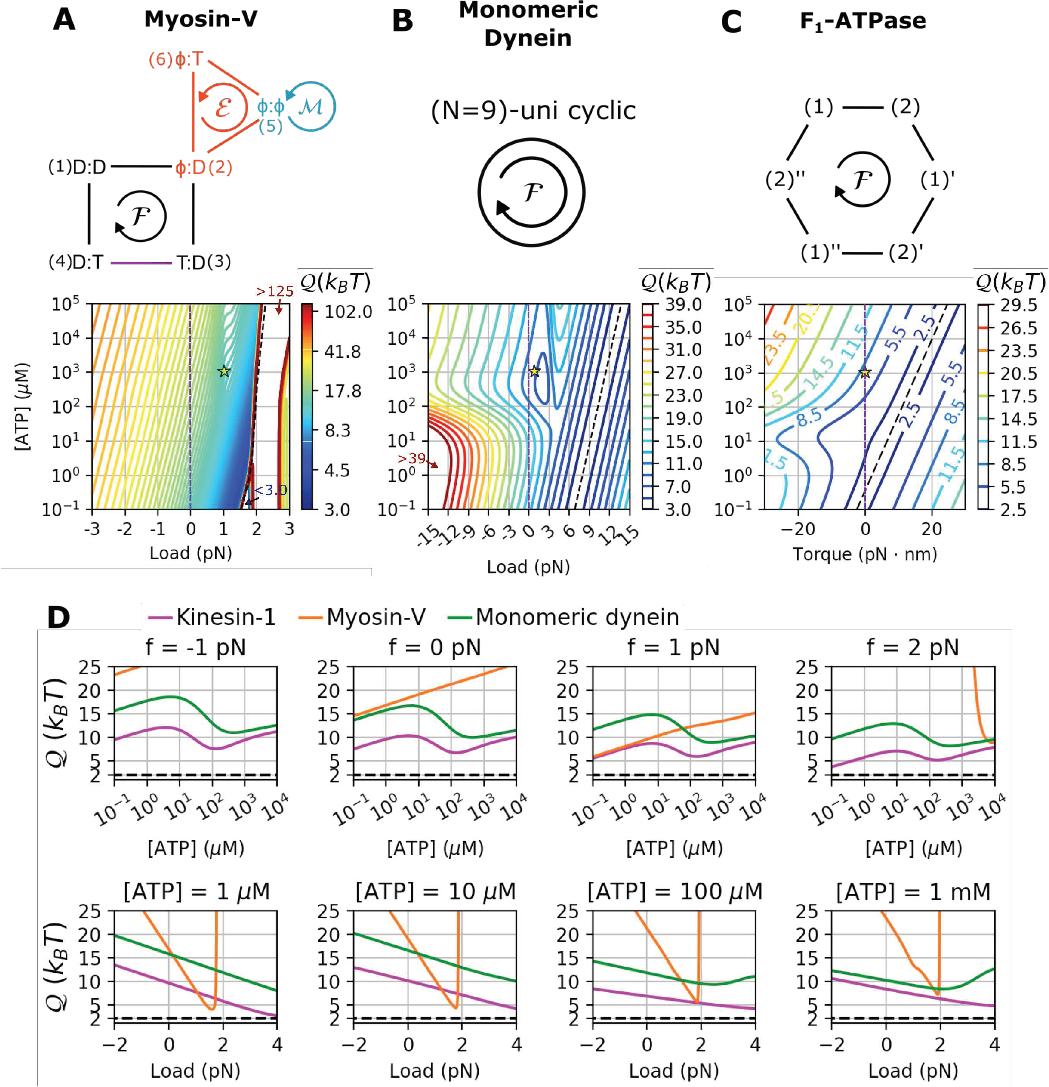
𝒬 for various motors as a function of *f* and [ATP]. **A.** Kinetic model for myosin-V consisting of three cycles 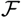, *ε*, and 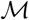 [38]. 𝒬(*f*, [ATP]) calculated at [ADP] = 70 *μ*M and [P_*i*_] = 1 mM (See also Fig. S11D for 2-D heat map). **B.** Kinetic model for monomeric dynein [39], and the corresponding 𝒬(*f*, [ATP]) calculated at [ADP] = 70 *μ*M and [P_*i*_] = 1 mM (See also Fig.S13D for 2-D heat map). **C.** 𝒬(*f*, [ATP]) at [ADP] = 70 *μ*M and [P_*i*_] = 1 mM (See also Fig.S13D for 2-D heat map) using the kinetic model for F_1_-ATPase from Ref. [40]. Other quantities such as *V*, *D*, and 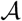 as a function of *f* and [ATP] are provided in Figs. S11, S13, S14. **D.** 𝒬(*f*, [ATP]) at fixed *f* (upper panels) and 𝒬(*f*) at fixed [ATP] (lower panels) for kinesin-1 (magenta), myosin-V (orange), and monomeric dynein (green).

**TABLE I.** Optimal *f* and [ATP] that locally minimize 𝒬. For unicyclic models of kinesin-1, myosin-V, and F_1_-ATPase, also shown are Δ𝒬 = 𝒬 − 𝒬_b_, where *Q*_b_ is a stronger lower bound for unicyclic kinetic schemes [5] (Eq.S40).

| | Kinesin-1<br>(multi-cycle) | Kinesin-1<br>(unicycle) | KIF17<br>(multi-cycle) | Myosin-V<br>(multi-cycle)<br>[ADP]=[P<sub>i</sub>]=0.1 μM | Myosin-V<br>(unicycle) | Dynein monomer<br>(unicycle) | F<sub>1</sub>-ATPase<br>(unicyclic) |
| --- | --- | --- | --- | --- | --- | --- | --- |
| $f$ (pN) | 4.1 | 3.2 <sup>a</sup> | 1.5 | 1.1 | 0.03 <sup>a</sup> | 2.1 | 8.6 <sup>a b</sup> |
| $[ATP]$ (μM) | 210 | 460 <sup>a</sup> | 200 | 20 | 17 <sup>a</sup> | 420 | 16 <sup>a</sup> |
| $\mathcal{Q}_{\min}$ ($k_B T$) | 4.0 | 4.5 | 9.2 | 6.5 | 14 | 8.1 | 4.2 |
| $\Delta\mathcal{Q}_{\min}$ ($k_B T$) | n/a | 1.6 | n/a | n/a | 0 | 5.1 | 0 |

<sup>a</sup> Condition for local minimization of $\Delta\mathcal{Q} = \mathcal{Q} - \mathcal{Q}_b$ .
<sup>b</sup> For F<sub>1</sub>-ATPase, we consider a resisting torque ( $\tau$ with the unit of pN·nm) against the rotation of the motor.

#### Dynein

𝒬 for monomeric dynein evaluated using the (N=9)-unicyclic model in Ref. [39] (Fig. 4B) is locally minimized to 𝒬 ≈ 8.1 *k*_*B*_*T* at *f* = 2.1 pN, [ATP] = 420 *μ*M (Fig. 4D, S13D). Compared to kinesin-1,𝒬 for monomeric dynein is optimized at lower *f*.

#### F_1_-ATPase

𝒬 for F_1_-ATPase using the (N=6)-state unicyclic model in Ref. [40] reveals that there is a valley around torque *τ* ≈ −10 pN·nm and [ATP] ≈ 10 *μ*M reaching 𝒬 ≈ 4 *k*_*B*_*T* (Fig. 4C). Notably, 𝒬 for F_1_-ATPase is optimized at *τ* < 0 in which ATP is synthesized, which is consistent with biologically expected role of F_1_-ATPase as an ATP synthase *in vivo.*

To compare between the motors, we plot 𝒬(*f*, [ATP]) at fixed *f* and 𝒬(*f*) at fixed [ATP] in Fig. 4D. 𝒬_kinesin-1_ < 𝒬_monomeric dynein_ < 𝒬_myosin-V_ over the broad range of *f* and [ATP]. We note that 𝒬_myosin-V_ is smaller than other motors at low [ATP] (= 1−10 *μ*M) and at small *f* (=1−2 pN).

## CONCLUDING REMARKS

Biological motors are far superior to macroscopic machines in converting chemical free energy into linear movement. The thermal noise is utilized to rectify the ATP binding/hydrolysis-coupled conformational dynamics into unidirectional movement, which conceptualizes the Brownian ratchet [42], but it also comes with a cost of overcoming the thermal noise that makes the movement of biological motors inherently stochastic and error-prone. Mechanism of harnessing energy into faster and more precise motion is critical for the accuracy of cellular computation. The uncertainty measure 𝒬 assesses the efficiency of improving the speed and regularity of dynamics for a given energetic cost.

Here, we have quantified the efficiency of biological motors using the uncertainty measure 𝒬 and found that the efficiencies of motors in cargo transport are all semi-optimized near the cellular condition (star symbols marking f ≈ 1 pN and [ATP]= 1 mM in Figs.2, 3, 4). *Q* versus *∊*_*l*_ plots (Fig.5, *∊*_*θ*_ for F_1_-ATPase motor) for the various motors, sorting the biological motors in the increasing order of 𝒬, are reminiscent of the recent study on the free-energy cost of accurate biochemical oscillations [43]. The plots indicate that kinesin-1 is the best motor whose 𝒬(≈ 7.2 *k*_*B*_*T*) approaches the bound of ideal motor (𝒬 = 2 *k*_*B*_*T*). Note that 𝒬(≈ 19 *k*_*B*_*T*) for the mutant kinesin-1, Kin6AA, is significantly greater than that for kinesin-1. The structure of 𝒬(*f*, [ATP]) and the suboptimal condition of 𝒬 differ from one type of motor to another. Here, we note that the structure of 𝒬(*f*, [ATP]) differs significantly from that of other quantities such as the flux *J*(*f*, [ATP]) [44] (equivalent to *V*(*f*, [ATP]) and the power efficiency *η*(*f*, [ATP]) ≡ *Ẇ*(*f*, [ATP])/*Ė*(*f*, [ATP]) (see **SI** Figures). - For example, compare 𝒬(*f*, [ATP]) (Fig.2A), *V*(*f*, [ATP]) (Fig.1D), and *η*(*f*, [ATP]) of kinesin-1 (Fig.5B) - Remarkably, it is 𝒬, not *V* nor *η*, that is effectively optimized over the range of *f* and [ATP] around the cellular condition. Further, as seen in Kin6AA, a modification of structure not only disturbs such suboptimality but also increase 𝒬 dramatically.

**FIG. 5.**
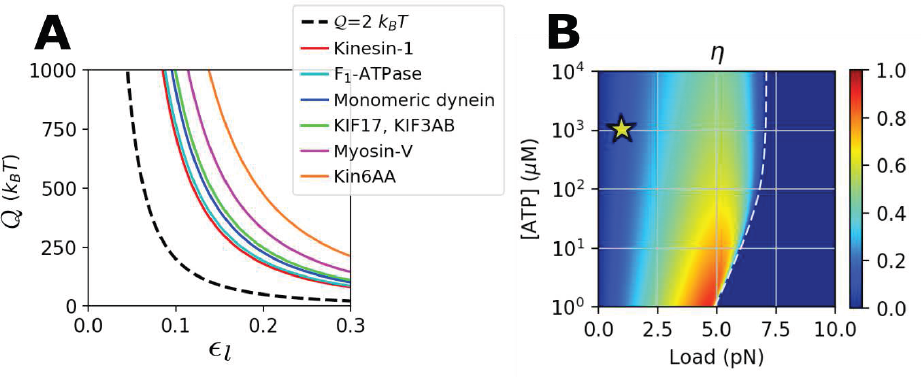
**A**. Plots of *Q* versus *∊*_*i*_ for various motors in cargo transport under the cellular condition, [ATP] = 1 mM [45] and *f* =1 pN [46, 47]. The plot for the ideal motor (𝒬 = 2*k*_*B*_*T*) is plotted with a dashed line. 𝒬/*k*_*B*_*T* = 7.2 (kinesin-1), 7.7 (F_1_-ATPase), 9.1 (monomeric dynein), 9.9 (KIF17), 9.9 (KIF3AB), 13 (myosin-V), 19 (Kin6AA). Zero torque is assumed for F_1_-ATPase. **B**. Power efficiency of kinesin-1 calculated using 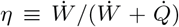. For *f* > *f*_stall_, the motor moves backward and *Ẇ* < 0. Thus, we set *η* = 0 for convenience. The star symbol indicates *f* ≈ 1 pN and [ATP] ≈ 1 mM.

To what extent can our findings on the *in vitro* single motor properties be generalized into those in live cells? First, the force hindering the motor movement varies with cargo size and subcellular location; the load or viscoelastic drag exerted against motors inside the cell varies dynamically [48, 49]. Yet, actual forces opposing the cargo movement in cytosolic environment are ≲ 1 pN [46, 47]. Since 𝒬s for microtubule-binding motors, kinesin-1, kinesin-2, and dynein, are narrowly tuned, varying only a few *k*_*B*_*T* over the range of 0 ≤ *f* ≤ 4 pN at [ATP] = 1 mM (Figs. 3D, 4D), our discussion on the *in vitro* single motor property in terms of 𝒬 can be extended to cargo transport in cytosolic environment. Next, a team of motors is often responsible for cargo transport in the cell [50]. It has, however, been shown that *in vitro* two kinesin motors attached to a cargo do not coordinate under low load, and saturating ATP [51]. Although trajectories generated by multiple motors have not been analyzed here, extension of the present analysis to such cases should be straightforward.

In axonal transport, of particular importance is the timely and efficient delivery of cellular material, the failure of which is linked to neuropathology [52, 53]. Since there are already numerous regulatory control mechanisms as well as other motors, it could be argued that the role played by the optimized single motor properties dictated by 𝒬 is redundant in light of the overall function of axonal transport. Yet, given that cellular regulations are realized through multiple layers of checkpoints [2], the optimized efficiency of motors at single molecule level can also be viewed as one of the checkpoints that assure the optimal cargo transport.

## METHODS

To study the transport properties of molecular motors, we used experimental data, available in the literature, of *V* and *D* under varying conditions of *f* and [ATP]. Once a set of kinetic rate constants {*k*_*ij*_} that defines the network model is determined by fitting the data of *V*({*k*_ij_(*f*, [ATP])}) and *D*({*k*_*ij*_(f, [ATP])}), it is straightforward to calculate 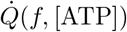 [12], and hence 𝒬(*f*, [ATP]).

For kinesin-1, we considered a 6-state double-cycle kinetic network (Fig.1A) and modeled the load-dependent rate constants differently from other kinetic steps [15]. The load dependence of kinetic rate was modeled using 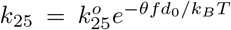 and 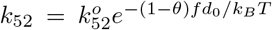 for the mechanical step between the states (2) and (5), and 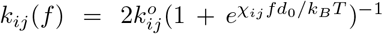 for other steps associated with ATP chemistry. The condition of *χ_ij_* = *χ_ji_* makes only the mechanical transition contribute to the work [15]. The rate constants determined in the 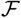-cycle are used for the corresponding chemical transition steps in the 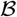-cycle [15]. For example, ADP dissociation rate constant *k*_23_ of 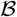-cycle is equal to *k*_56_ which describes ADP dissociation in 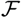-cycle. Similarly, *k*_32_ = *k*_65_, *k*_34_ = *k*_61_, *k*_43_ = *k*_16_,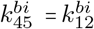, *χ*_23_ = *χ*_56_, *χ*_34_ = *χ*_61_, *χ*_45_ = *χ*_12_. *k*_54_ is determined from the equality (*k*_12_*k*_25_*k*_56_*k*_61_/*k*_21_*k*_52_*k*_65_*k*_16_) = (*k*_23_*k*_34_*k*_45_*k*_52_/*k*_32_*k*_43_*k*_54_*k*_25_) which results from the fact that ATP hydrolysis free energy that drives the 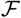- and 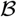-cycle is identical, and hence *k*_54_ = *k*_21_(*k*_52_/*k*_25_)^2^ [15]. As the experimental data from Ref. [31] are scarce at high load condition, which activates the 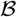-cycle, it is not easy to determine all the parameters for 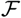 and 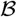 cycles simultaneously using the existing data. To circumvent this difficulty, we fit the data using the following procedure. First, the affinity 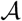 at *f* = 0 was determined from our previous study that employed the (N=4)-state uni-cyclic model [12]. Even though 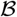-cycle is not considered in Ref. [12], 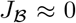 at *f* ~ 0, which justifies the use of unicyclic model at *f* ≪ *f*_stall_. Next, the range of parameters were constrained during the fitting procedure (Table III) based on the values obtained in [15, 16]). To fit the data globally, we employed the `minimize` function with ‘L-BFGS-B’ method from the scipy library.

For Kin6AA, KIF17, and KIF3AB, motility data digitized from Ref. [18, 32] were fit to the same 6-state network model used for kinesin-1. For myosin-V, dynein, and F1-ATPase, we employed kinetic network models and corresponding rate constants used in Ref. [38-40]. Further details are provided in **SI**.

## ACKNOWLEDGEMENTS

We thank Steven P. Gross for insightful comments on cargo transport *in vivo*. This work was performed in part at Aspen Center for Physics, which is supported by National Science Foundation grant PHY-1607611. We acknowledge the Center for Advanced Computation in KIAS for providing computing resources.

## Supplementary Information

### CALCULATION OF *V* AND *D* OF KINESIN-1 IN 6-STATE MULTI-CYCLIC MODEL

To obtain the expression of *V* and *D* for multicyclic kinetic network model in terms of a set of rate constants [*k*_*ij*_}, we have generalized the technique by Koza [13] (Alternatively, technique based on the large deviation theory can be used. See Ref. [14, 54]). We define the generating functions for the given network model.

In the 6-state double-cycle kinetic network (Fig. 1A), we define the three distinct generating functions for 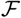, 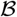, and *χ* cycles. The two generating functions for the subcycles, 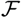 and 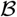-cycles, are convenient to calculate the chemical current 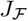 and 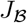 in each subcycle. To calculate *V* and *D* in a convenient way, we have defined another generating function for *χ*-cycle, which is not explicit in the kinetic scheme in Fig. 1A. The *χ*-cycle differs from 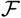, 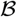-cycle in that the former explicitly considers the physical location of the motor along the 1D track. Although *V* is obtained either evaluating 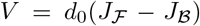 or *V* = *d*_0_*J*_*χ*_, it is not straightforward to decompose the diffusivity of motor *D* into the contributions from 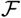 and 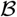-cycle.

The expressions of 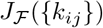, 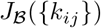, *V*([*k*_*ij*_}), and *D*([*k*_*ij*_}) can be obtained by considering an asymptotic limit (*t* → ∞) of the corresponding generating function.

In what follows, we provide the derivation of generating function in details. In order to derive the generating function, we introduce a generalized index for reaction cycle 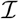, with 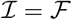, 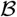, or 𝒳.

### Master equation

For a system with *N* chemical states ([1, 2, ⋯, *N*}), a generalized state 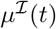 is defined by using the chemical state of the motor at time *t* and the number of completed 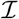-cycles 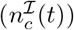. For kinesins whose dynamics can be mapped onto the 6-state double-cycle kinetic network model, if the motor is in the *i*-th chemical state (*i* ∈ [1, 2, ⋯, *N*} with *N* = 6) at time *t*, the generalized state of the motor in the 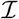-cycle is 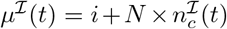, where 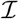 could denote either 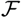, 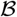, or 𝒳 depending on reader’s interest. 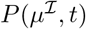 that represents the probability of the system being in 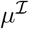 at time *t*, satisfies

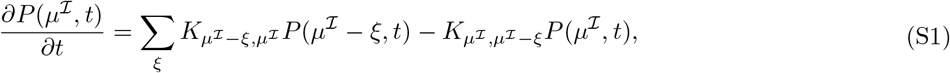

where 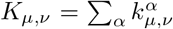 and 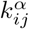 denotes the rate of transition from state *i* to state *j* that follows the *α*-th pathway. Here, the periodicity of network model imposes 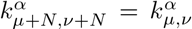, *K*_*v*+*N*, *μ*+*N*_ = *K*_*v*,*μ*_, and 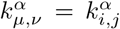 for *μ* = *i* (mod N) and *v* = *j* (mod N). The range of (integer) summation index *ξ* depends on the existing path-ways for 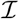-cycle. Hereafter, the superscript 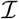 on *μ* shall be omitted for simplicity.

Following Ref.[13],we define

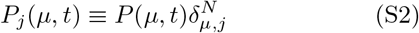

where,

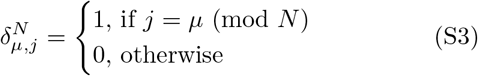

Here, *j* ∊ [1, 2, ⋯, *N*}. Multiplying on 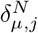 both sides of Eq.S1 and using the equality 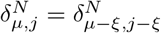, we get

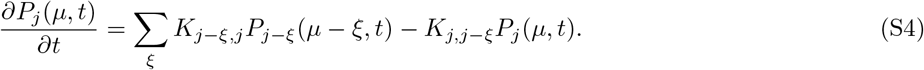

### Generating function

We define a generating function to derive *V* and *D*. The generating function for 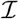-cycle is defined by

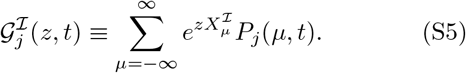

where 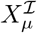 denotes the generalized coordinate for 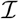-cycle at generalized state *μ*. Then Eq.(S4) and the equality 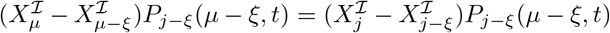 with 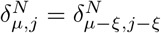 lead to

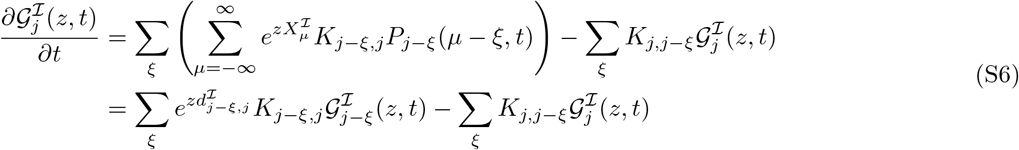

where 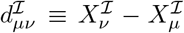. In general, different cycle has different 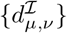. For example, for the 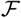-cycle in Fig. 1A,

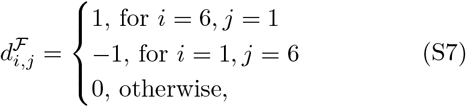

for the 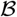-cycle,

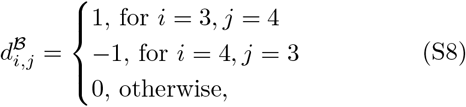

and for the 𝒳-cycle,

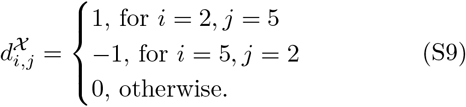

In fact, Eq. (S6) can be expressed more succinctly as

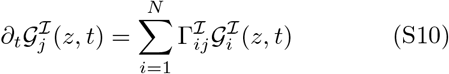

where

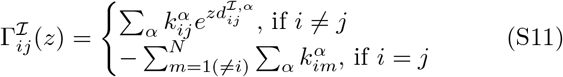

With *α*, an index to discern the pathways, Г can be written in the form of *N* × *N* matrix.

### Generating function at the asymptotic limit

Here we consider the asymptotic limit (*t* → ∞) in which *V* and *D* are well defined for an arbitrary chemical network model. The general solution of Eq.(S10) can be written as [13]

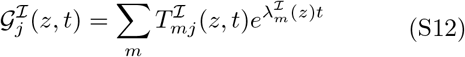

where 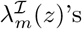 (*m* = 0, 1, 2, ⋯, *N*) are the eigenvalues of 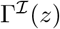. For a system in (unique) steady state, the eigenvalues satisfy 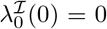 and 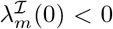 for *m* ≠ 0. Thus, at t → ∞ and when *z* ~ 0,

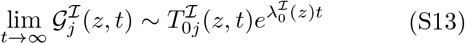

Now, summed over the index *j*, Eq.(S5) is led to

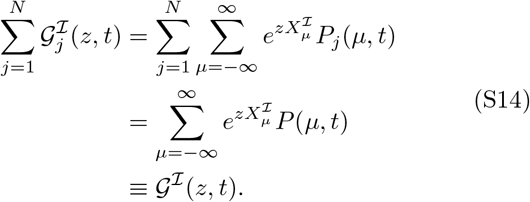

From Eq.(S13), at *t* → ∞ we have

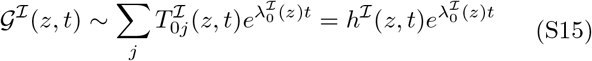

where 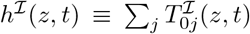. Since 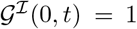 and 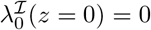, 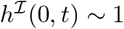 at *t* → ∞.

### Velocity and Diffusion coefficient

In this section, we first define the flux 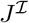 and the diffusion coefficient 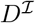 of 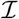-cycle using 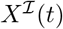 at *t* → ∞. Then by using the asymptotic form of the generating function, we will get the relation between 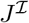 and 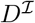, and the lowest eigenvalue 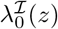.

The mean value of the generalized coordinate 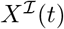 can be obtained using

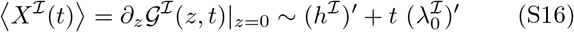

where Eq.(S15) was used and the prime denotes a partial derivative with respect to *z* at *z* = 0. The flux of 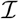-cycle is defined by

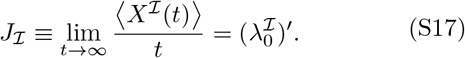

*J*_*χ*_ multiplied by the step size *d*_0_ corresponds to the velocity *V* of motor

Similarly, the diffusion coefficient 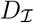 is obtained by considering the second moment of 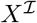.

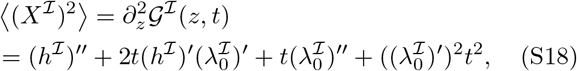

which gives

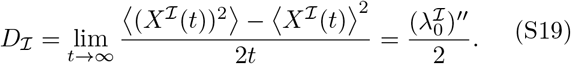

Thus, the diffusion coefficient of motor is obtained: 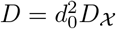.

### Characteristic polynomial

To express the derivatives of 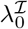 in terms of rates {*k*_*ij*_}, we use the characteristic polynomial of 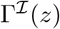 [13],

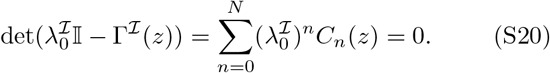

By differentiating both side of Eq.S20 with respect to *z* and setting *z* = 0, we get

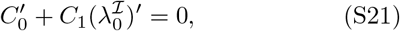

and

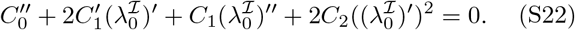

From Eqs.(S21) and (S22),we get

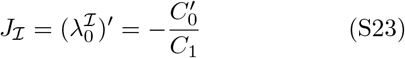

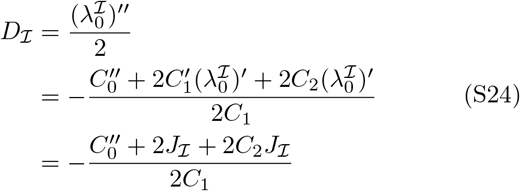

*C*_n_’s and their derivatives, which depend on the choice of 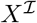, can readily be found by differentiating the characteristic polynomial with respect to 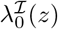 with 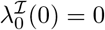 [13].

### Explicit expression of 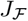

The expression of reaction current in each subcycle 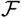 and 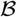 in terms of {*k*_*ij*_} can be obtained by considering the corresponding generating function 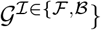 Here, we provide the expression of 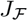 in terms of rate constants {*k_ij_*} for the 6-state double-cycle kinetic network.

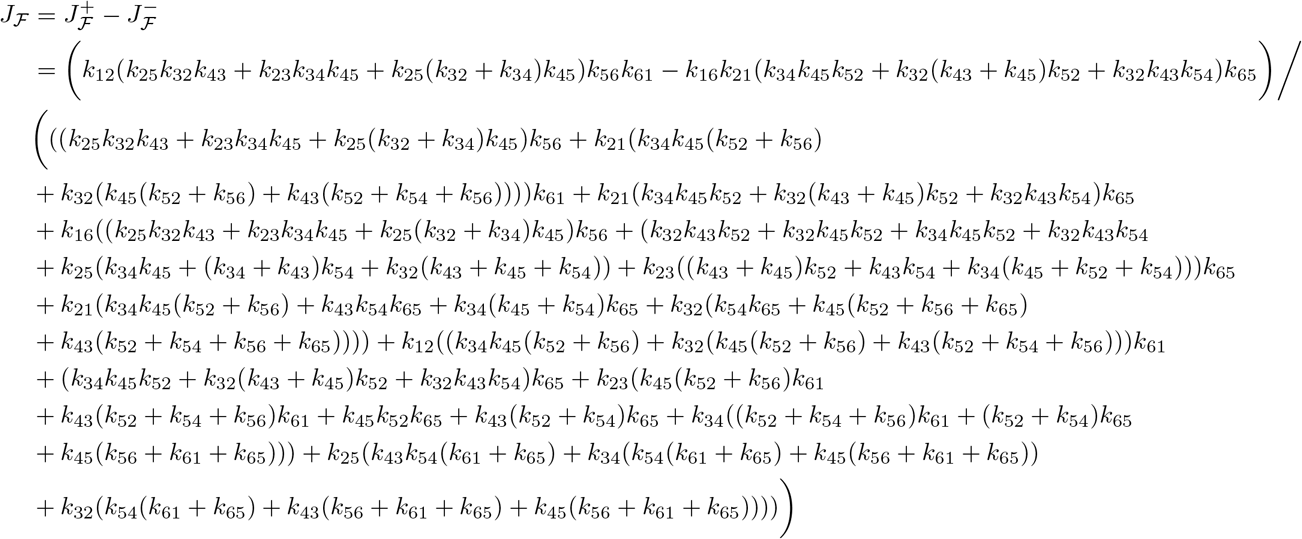

Similarly, 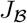 and *D*_*χ*_ can also be expressed in terms of {*k*_*ij*_}.

## ANALYSIS OF OTHER TYPES OF KINESINS

### Kinesin-1 mutant (Kin6AA)

Single molecule motility data digitized from Ref. [55] was fitted to 6-state network model (Figs. 1A, S4) by using the same method employed for the analysis of kinesin-1 data (**Methods**). However, 4 additional initial conditions for *k*_25_ ({300, 3000, 30000, 3000000}), thus total 245 initial conditions, were explored. The rate constants estimated from this procedure are provided in Table II.

**TABLE II.** Parameters determined for the 6-state double-cycle model [15]. The unit of rate constants ({*k*_*ij*_}) is *s*^−1^ except for 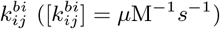. The rates in the table are determined for *f* = 0.

|  | Kinesin-1 | Kin6AA | KIF17 | KIF3AB |
| --- | --- | --- | --- | --- |
| $k_{12}^{bi}$ | 2.8 | 10 | 10 | 10 |
| $k_{21}$ | 4200 | 92 | 3600 | 500 |
| $k_{25}$ | $1.6 \times 10^6$ | $1.6 \times 10^4$ | $4.0 \times 10^4$ | $6.2 \times 10^6$ |
| $k_{52}$ | 1.1 | 3.4 | 0.079 | 7.2 |
| $k_{56}$ | 190 | 680 | 590 | 92 |
| $k_{65}$ | 10 | 4.1 | 13 | 37 |
| $k_{61}$ | 250 | 58 | 310 | 320 |
| $k_{16}$ | 230 | 260 | 1100 | 750 |
| $k_{54}$ | $2.1 \times 10^{-9}$ | $4.3 \times 10^{-6}$ | $1.4 \times 10^{-8}$ | $6.8 \times 10^{-10}$ |
| $\theta$ | 0.61 | 0.59 | 0.34 | 0.82 |
| $\chi_{12}$ | 0.15 | 0.12 | 0.15 | 0.09 |
| $\chi_{56}$ | 0.0015 | 0.0 | 0.012 | 0.021 |
| $\chi_{61}$ | 0.11 | 0.18 | 0.17 | 0.16 |

### Kinesin-2 (KIF17, KIF3AB)

Single molecule motility data digitized from Ref. [32] was again fitted to the 6-state double-cycle kinetic model (Fig. 1A, S6, S8) following the identical procedure employed in the analysis of kinesin-1 data (**Methods**). However, two additional initial conditions for k_25_ ({30000, 3000000}) were explored, which results in total 147 initial conditions. The rate constants are shown in Table II.

### MYOSIN-V

Here we summarize the multi-cyclic model for myosin-V [38] which consists of ATP-dependent chemomechanical forward cycle 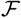, dissipative cycle *ε*, and ratcheting cycle (ATP independent stepping cycle) 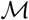 (Fig. 4A). We first provide the explanation of how *V* and *D* of myosin-V are calculated. Next, the affinity and heat production 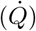 are expressed in terms of a set of rates {*k*_*ij*_}. Finally, 𝒬 shall be calculated using *V*, *D*, and 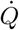.

### Calculation of *V* and *D*

The 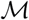-cycle consisting of a single state (Fig. 4A) prevents the application of Eq.(S11). To circumvent this difficulty, the model with additional state (5′) is considered (Fig. S10). The (5′)-state is chemically equivalent to the state (5), but describes motor in different position on actins, such that *X*(5) = *X*_0_ and *X*(5′) = *X*_0_ ± *d*_0_ where *d*_0_ = 36 nm for myosin-V. In this new network, the rate constants *κ_ij_*’s are

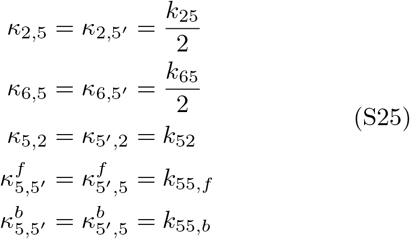

where the subscripts *f* and *b* denote the forward and backward motion, respectively. Other rate constants satisfy *κ*_*ij*_ = *k*_*ij*_. This modification can be justified by considering stochastic movement of myosin-V on the chemical network [56]: *κ*_*i*,5_, *ε*_*i*,5′_ are set to *k*_*i*5_/2, such that the outgoing fluxes from the states *i* = (2), (6) to the state (5) remain identical in the both networks depicted in Fig. 4A and Fig.S10. Next, we set *κ*_5,*i*_ = *k*_5′,*i*_ to keep the inward fluxes toward (6), (2) identical for the two networks. Finally, 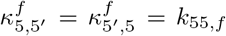 and 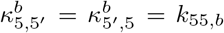. These modification of rate constants enable us to describe transitions within the 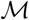-cycle.

Now, the elements of distance matrix scaled by *d*_0_ are

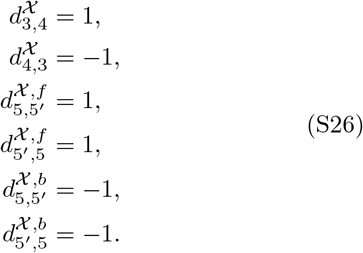

Other elements 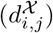 are all zero. Thus, 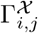 is written as (with (7) = (5′))

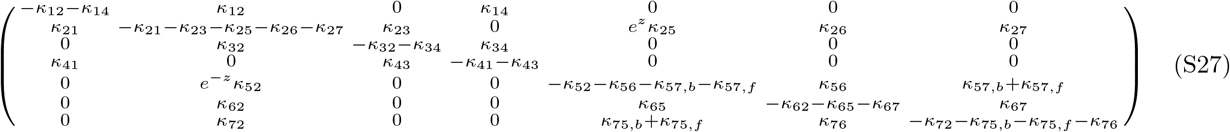

Now, the travel velocity *V* and the diffusion coefficient *D* of myosin-V are readily acquired by using Eqs.(S23, S24, and S27).

### Affinities and heat production

The affinities of individual cycles are

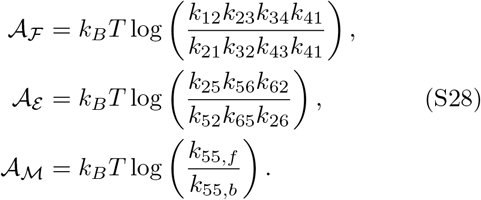

Only the following rate constants depend on the load (*f*):

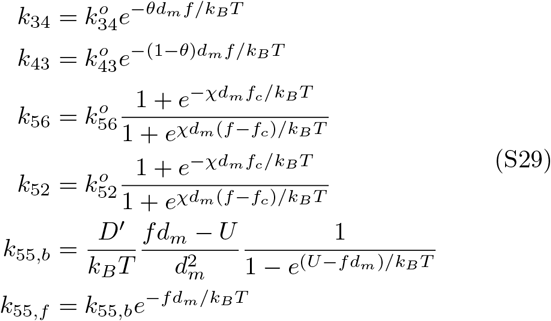

where

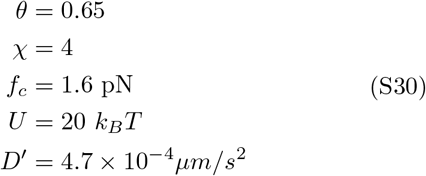

as described in Ref. [38]. Thus, the affinities can be written as

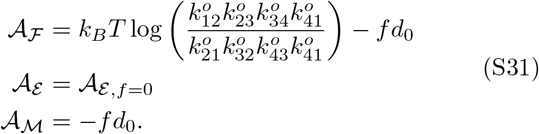

The relation 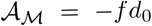 results from the fact that 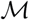-cycle is ATP-independent and activated by the load. Thus, the heat production rate of the system is

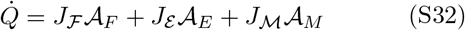

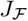, *J*_*ε*_, and 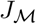 can be calculated by using Eqs. (S23) and (S27). Finally, 𝒬 for myosin-V is given by

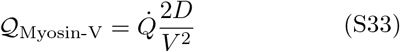

where 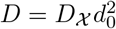 and *V* = *J*_*χ*_*d*_0_.

### MONOMERIC DYNEIN

The unicyclic model with 9 chemical states for the movement of monomeric dynein is considered based on the model shown in Fig. 3 of Ref. [39]. Identical rates constants shown in Table 3 of Ref. [39] were employed for the calculation. To describe force-dependence, we model the rate constant for power stroke *k*_+*PS*_ and for reverse stroke *k*_−*PS*_ as follows.

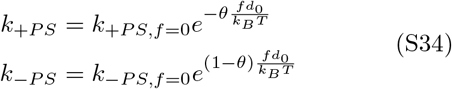

where *θ* = 0.3 is selected based on the previous study [57, 58]. Again, *V* and *D* were calculated using Eqs. S23, S24.

**TABLE III.** Initial values and constraints applied during the fit of kinesin data using 6-state double-cyclic model. The units are identical to those in Table II. For *k*_65_ and *k*_16_, we used 7 initial values (0.001, 0.01, 0.1,1,10,100,1000) for the fits.

|  |  |  |  |  |  |  |  |
| --- | --- | --- | --- | --- | --- | --- | --- |
| $k_{12}^{bi}$ | $0.5 \leq 1.8 \leq 10$ | $k_{56}$ | $10 \leq 200 \leq 10^4$ | $k_{61}$ | $10 \leq 200 \leq 10^4$ | $k_{25}$ | $10^4 \leq 3 \times 10^5 \leq 10^7$ |
| $k_{21}$ | $10 \leq 100 \leq 10^4$ | $k_{65}$ | $10^{-4} \leq 10^{[-3, -2, -1, 0, 1, 2, 3]} \leq 10^4$ | $k_{16}$ | $0^{-4} \leq 10^{[-3, -2, -1, 0, 1, 2, 3]} \leq 10^4$ | | |
| $\theta$ | $0 \leq 0.3 \leq 1$ | $\chi_{12}$ | $0 \leq 0.25 \leq 1$ | $\chi_{56}$ | $0 \leq 0.05 \leq 1$ | $\chi_{61}$ | $0 \leq 0.05 \leq 1$ |

### Affinity and heat production

The affinity for unicyclic model is written as [12, 23, 59]

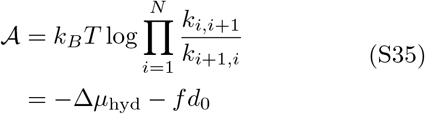

where *d*_0_ = 8.2 nm. The second term, describing force-dependence, is originated from the use of Eq. S34. Finally, 𝒬 is

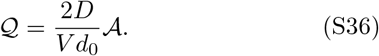

### F_1_-ATPASE

Here, we summarize the unicyclic model developed for F_1_-ATPase in Ref. [40]. The model is (*N* = 2) uni-cyclic model (Fig. 4C) where 3 cycles in chemical state space correspond to a single rotation in real space (angle changes by 90° upon transition from the state (1) to state (2) whereas transitions from the state (2) to (1)′ induce 30° rotation (Fig. 4C). The model is valid when the torque applied to F_1_-ATPase is small enough (*τ* ≲ 30 pN·nm) that the mechanical cycle is tightly coupled to the chemical reaction [40]. The dependences of rate constants on the torque are

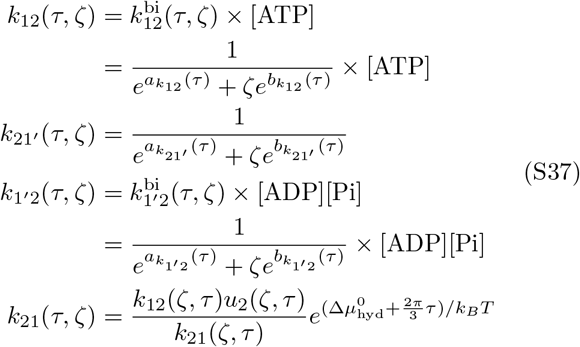

where 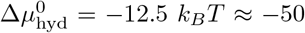, ζ is the friction coefficient (for example, if the γ-shaft of F_1_-ATPase is attached to a bead of radius *r*, ζ = 2π*ηr*^3^(4+3 sin^2^ π/6) [40] with the water viscosity *η* = 1 cP = 10^−9^ pN×s×nm^−2^. In our calculation, *r* = 40 nm as in Ref. [40]), and *a*_i_(*τ*),*b*_*i*_(*τ*) are polynomial function of *τ* defined in Ref. [40].

### *V*, *D*, affinities, and heat production

For (N=2)-unicyclic model, the speed of rotation *V*, diffusion coefficient *D*, and affinity 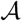 are [12, 13, 22, 23, 26, 60, 61]

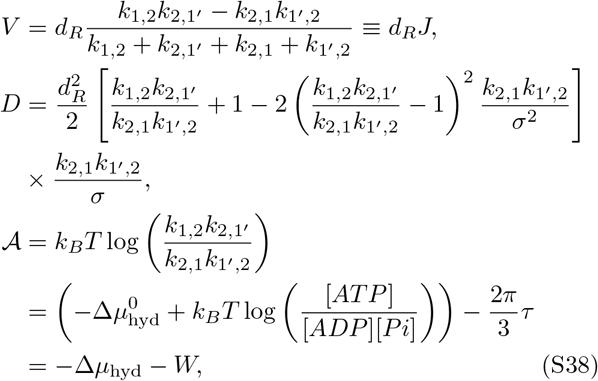

where 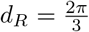 is the radian distance that motor travels upon ATP hydrolysis, *σ* = *k*_12_ + *k*_21′_ + *k*_21_ + *k*_1′2_, and 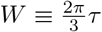 denotes the work done by the motor. Here, *τ* > 0 implies the motor performs work against the hindering load. Thus, 𝒬 is given by

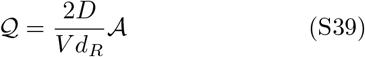

### UNICYCLIC KINETIC MODEL FOR KINESIN-1

To analyze the kinesin-1 data, we also considered (*N* = 4)-unicyclic model (Fig. S3A) which was used in our previous study [12]. Briefly, the model consists of four forward rates {*u*_1_, *u*_2_, *u*_3_, *u*_4_} and four backward rates {*w*_1_, *w*_2_, *w*_3_, *w*_4_}. Only *u*_1_(= *k*^*bi*^[ATP]) depends on [ATP]. Barometric dependence of the rates on forces are assumed: 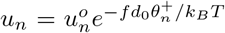 and 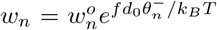 with 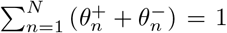 [27, 62]. *V*, *D*, *A*, and a set of rate constants used in the calculation of 𝒬 are available in Ref. [12]

### UNICYCLIC KINETIC MODEL FOR MYOSIN-V

For myosin-V, we also considered the (*N* = 2) uni-cyclic model from Ref. [41]. Briefly, the model consists of two forward rates {*u*_1_,*u*_2_} and two backward rates {*w*_1_,*w*_2_}. Only *u*_1_(= *k*[ATP]) and *w*_2_(= k′[ATP]^α^) depend on [ATP]. Here, *α* = 1/2. Different choice of *α* introduces only minor difference in the results as argued in [41]. Barometric dependences of the rates on forces are assumed again: 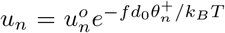 and 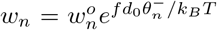 with 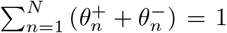 [27, 62]. The parameters used in the calculation are available in Eqs. (12), (13) in Ref. [41]. Identical expressions for *V*, *D*, and 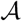 from Eq. S38 were used for the calculation except for *W* = *fd*_0_.

### THE LOWER BOUND OF 𝒬 FOR UNICYCLIC MODEL

The analytic expression for the lower bound of the uncertainty measure 𝒬 is available for unicyclic models [5]. For (*N*)-state unicyclic model, the lower bound of 𝒬 is

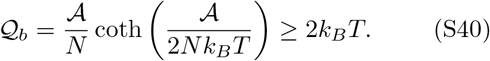

The 𝒬_*b*_ and the Δ𝒬 ≡ 𝒬−𝒬_*b*_ of the motors as a function of *f* and [ATP] are calculated in Figs. S3D (kinesin-1), S14D (F_1_-ATPase), and S15D (myosin-V).

**FIG. S1.**
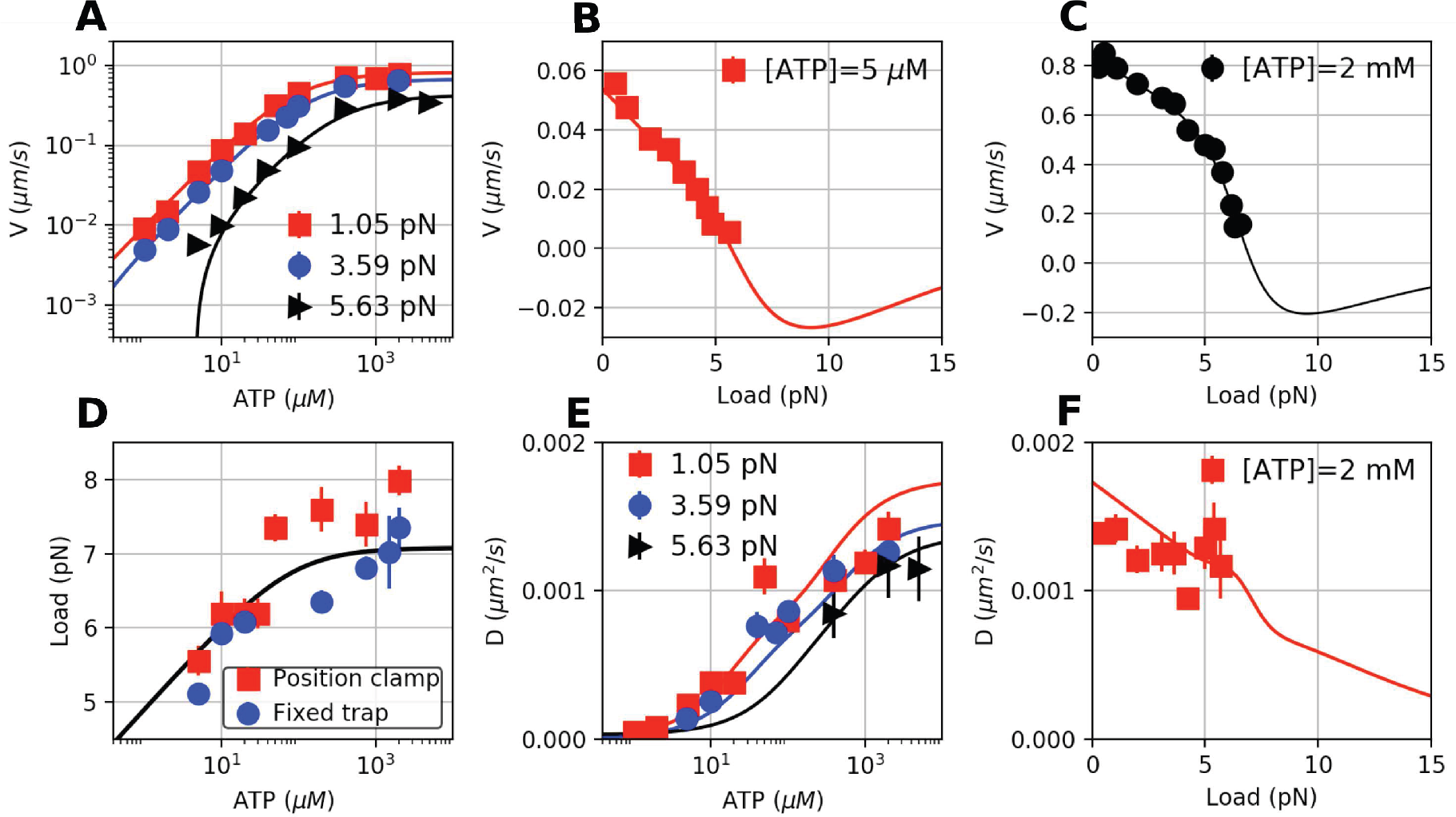
Analysis of experimental data of kinesin-1, digitized from Ref. [31], using the 6-state model [15]. The solid lines are the fits to the data **A.** *V* vs [ATP] at *f* =1.05 pN (red square), 3.59 pN (blue circle), and 5.63 pN (black triangle). **B.** *V* vs *f* at [ATP] = 5 *μ*M. **C.** *V* vs *f* at [ATP] = 2 mM. **D.** Stall force as a function of [AtP], measured by ‘Position clamp’ (red square) or ‘Fixed trap’ (blue circle) methods. **E.** *D* vs [ATP] at *f* =1.05 pN (red square), 3.59 pN (blue circle), and 5.63 pN (black triangle). *D* was estimated from *r* = 2*D*/*Vd*_0_. **F.** *D* vs *f* at [ATP] = 2 mM.

**FIG. S2.**
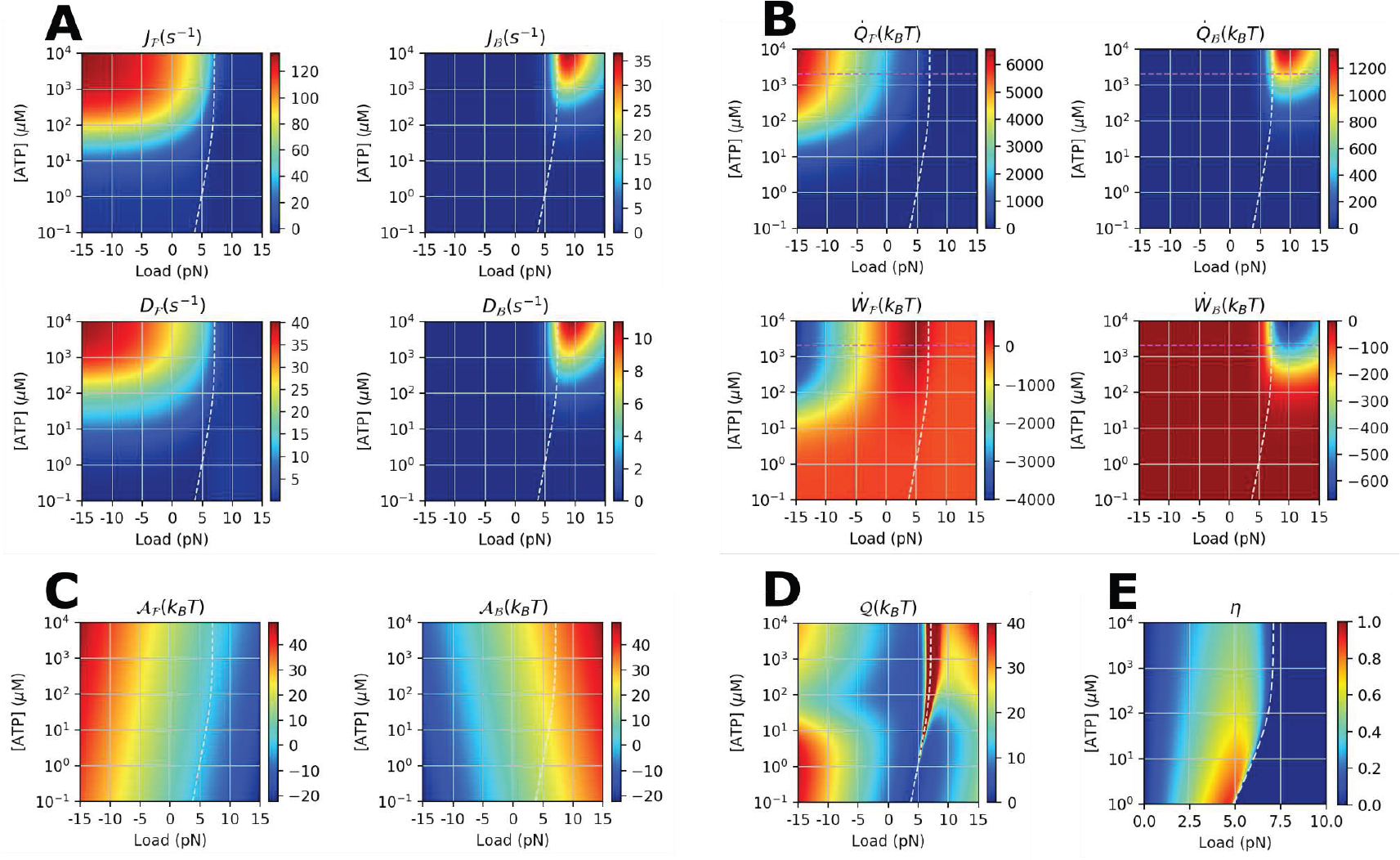
Various physical properties of kinesin-1 calculated using the 6-state network model (Fig. 1A) at varying *f* and [ATP]. **A.** Transport properties (flux *J* and *D*). **B.** Heat 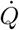 and work production *Ẇ*. **C.** Thermodynamic affinity 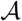. **D.** 𝒬(*f*, [ATP]). **E.** Power efficiency 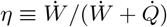 For *f* > *f*_stall_, *η* is set to zero. The white dashed lines indicate the stall condition.

**FIG. S3.**
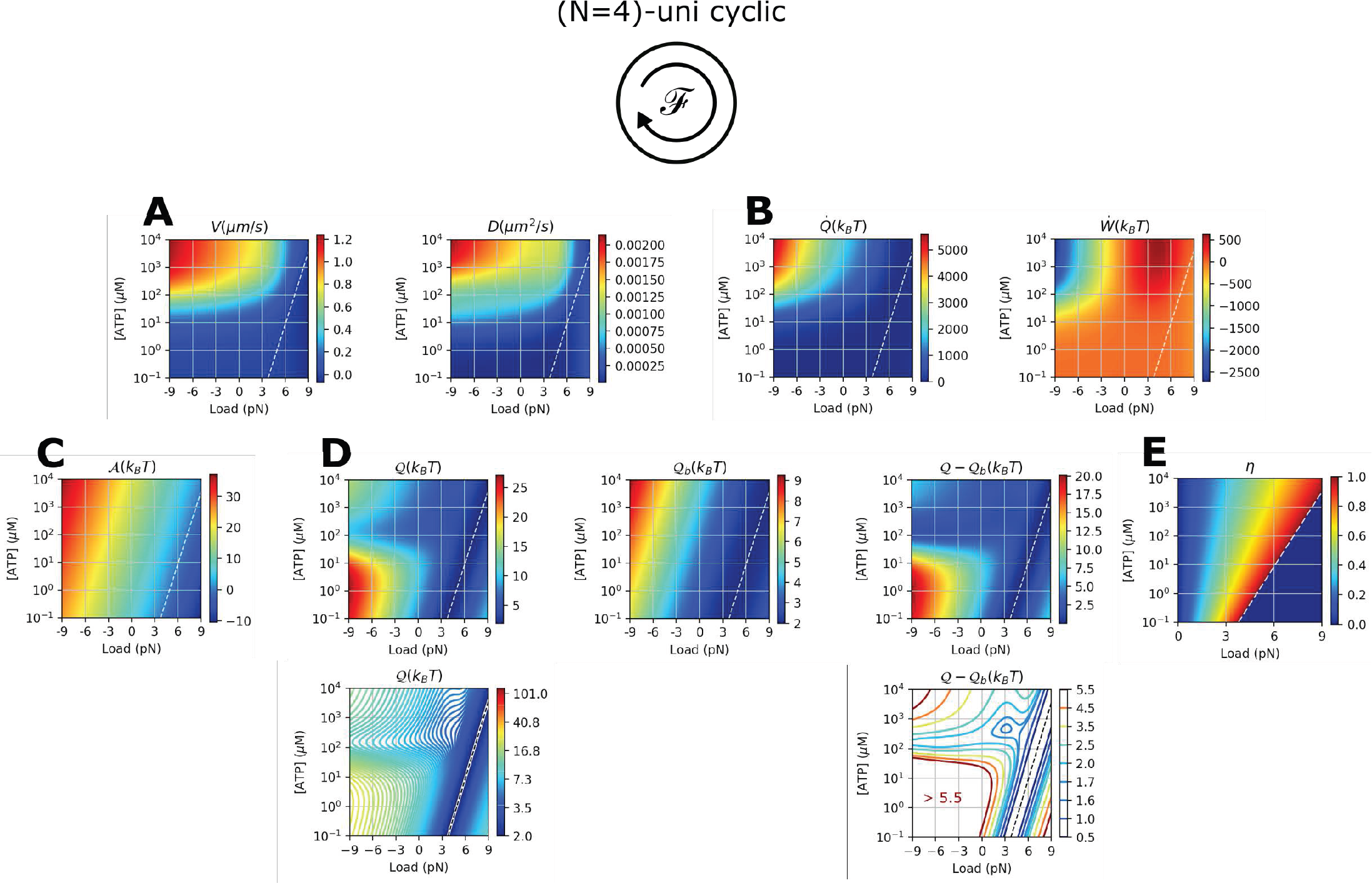
Various physical properties of kinesin-1 calculated using (N=4)-unicycle model [12, 27] at varying *f* and [ATP]. We used the same model parameters from Ref. [12]. **A.** Transport properties (*V*, *D*). **B.** Heat 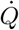 and work production *Ẇ*. **C.** Thermodynamic affinity 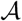. **D.** 𝒬, 𝒬_*b*_ (Eq.S40), and their difference Δ𝒬 = 𝒬−𝒬_*b*_. For clarity, identical data of 𝒬 and Δ𝒬 are shown again using contour plots. **E.** Power efficiency 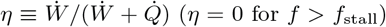. The white dashed lines indicate the stall condition.

**FIG. S4.**
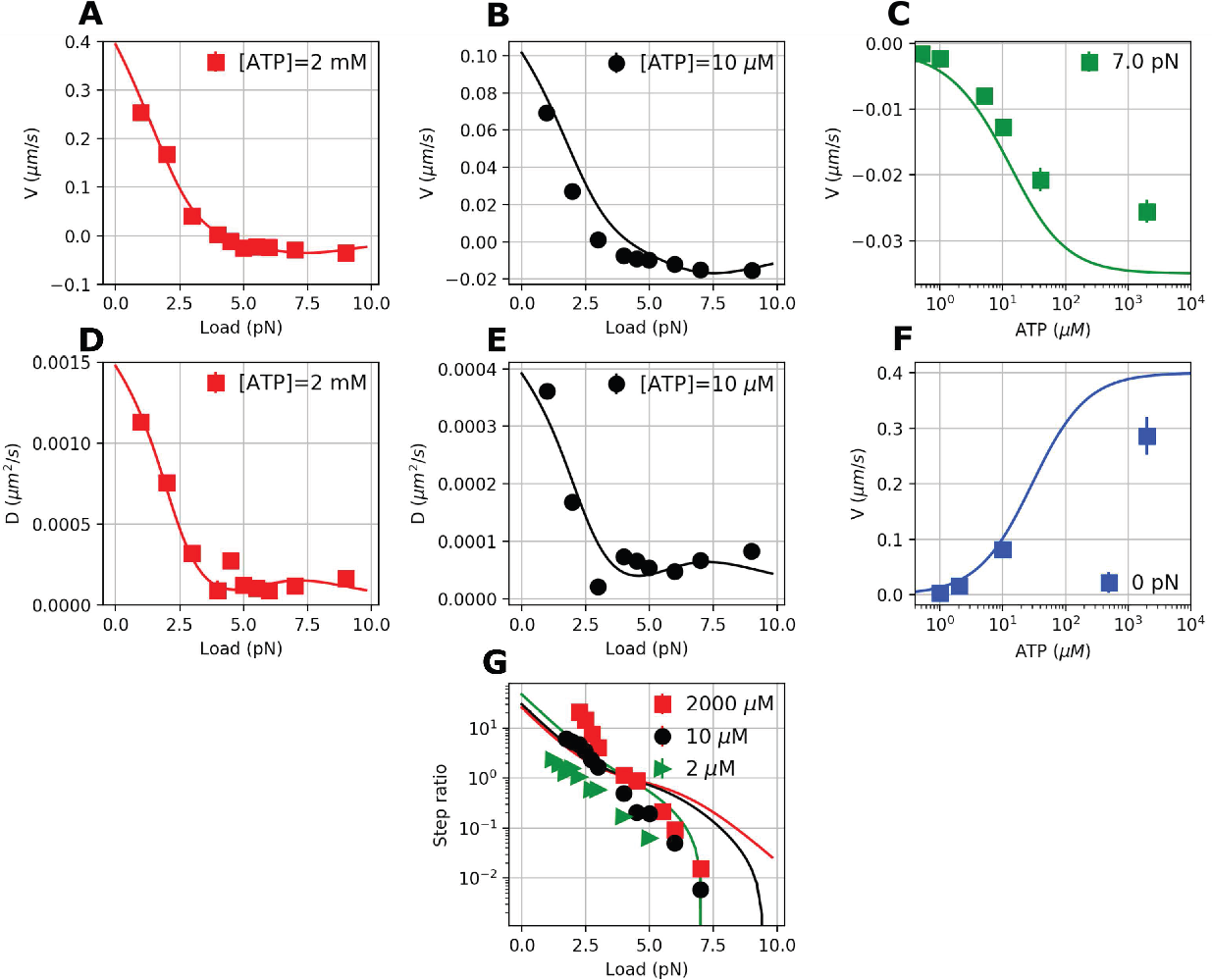
Motility data of Kin6AA, a mutant made of kinesin-1 to which six additional amino-acids are inserted in the neck-linker domains [55], and the theoretical fits made using the 6-state double-cycle kinetic network model (Fig. 1A).

**FIG. S5.**
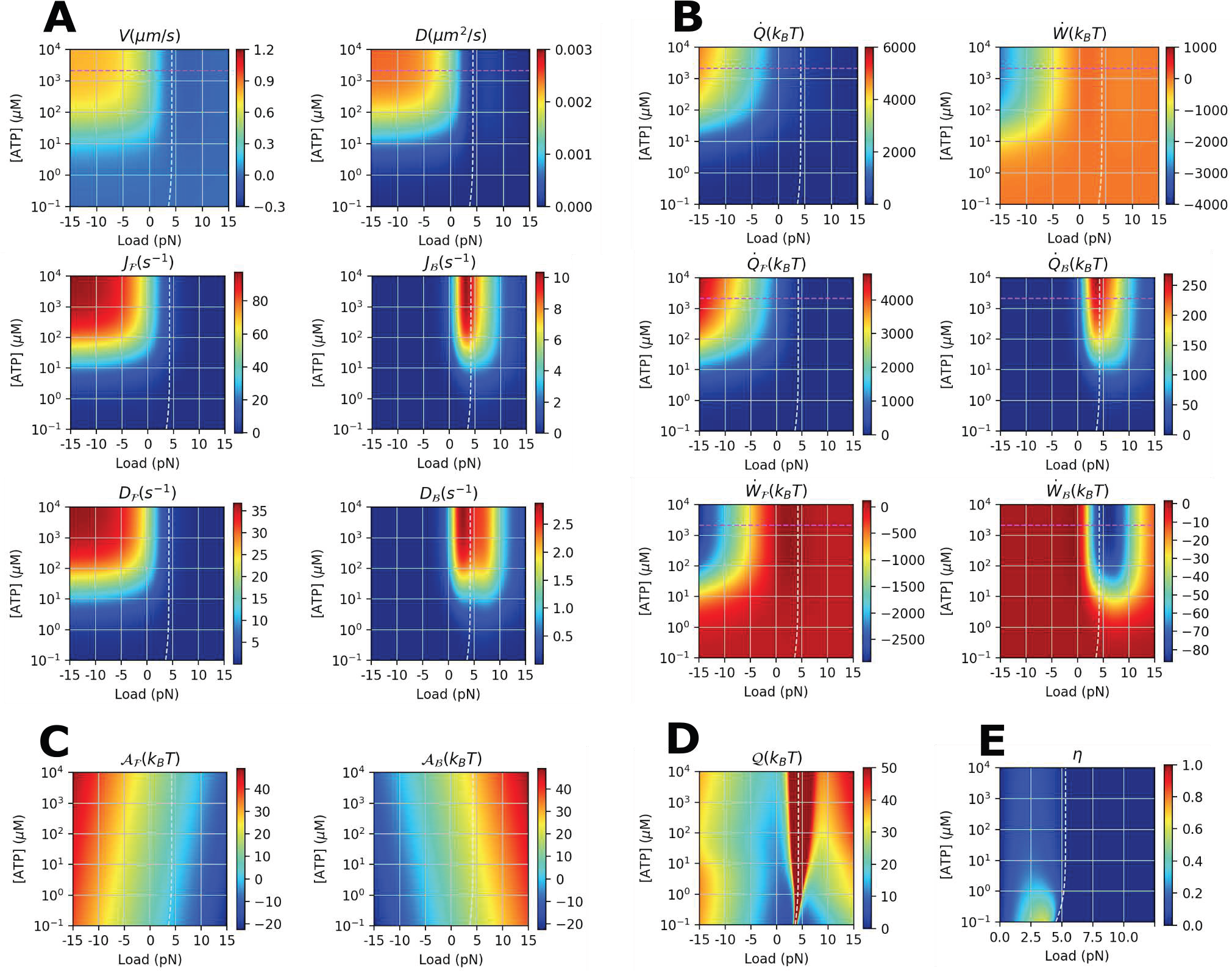
Various physical properties of kin6AA calculated using 6-state network model (Fig. 1A) at varying *f* and [ATP]. **A.** Transport properties (*J*, *D*). **B.** Heat 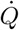 and work production *Ẇ*. **C.** Thermodynamic affinity 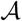. **D.** 𝒬(*f*, [ATP]). **E.** Power efficiency 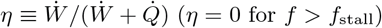. The white dashed lines indicate the stall condition.

**FIG. S6.**
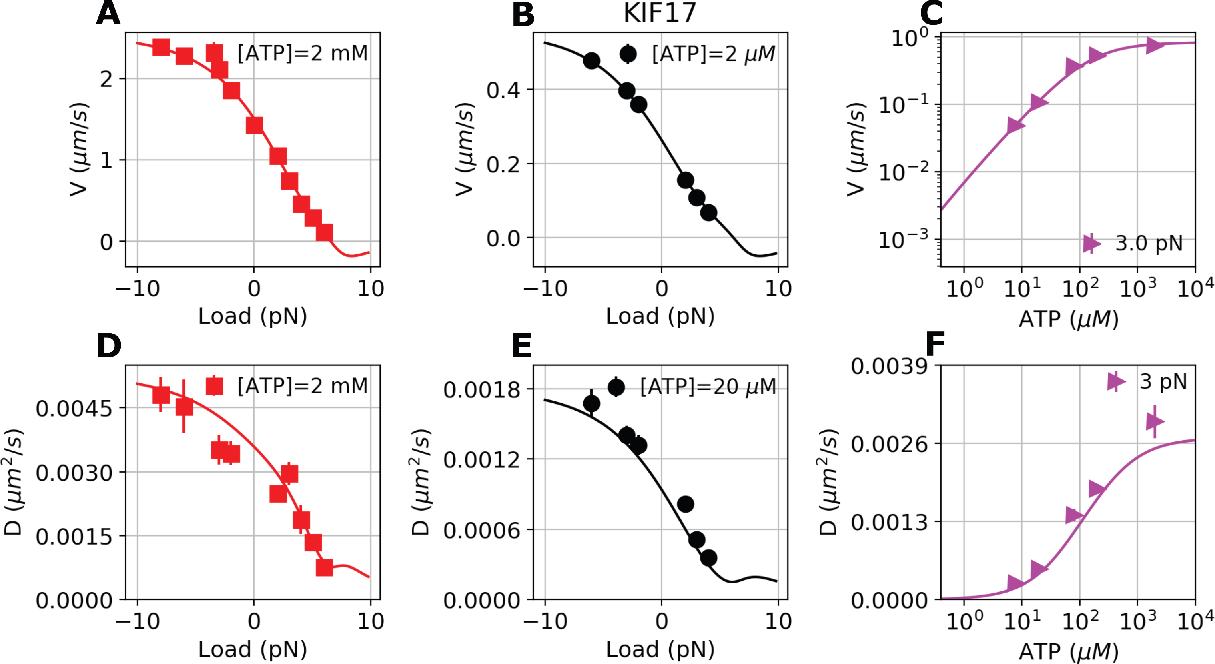
Motility data of homodimeric kinesin-2 (KIF17) [32] and their theoretical fits (solid lines).

**FIG. S7.**
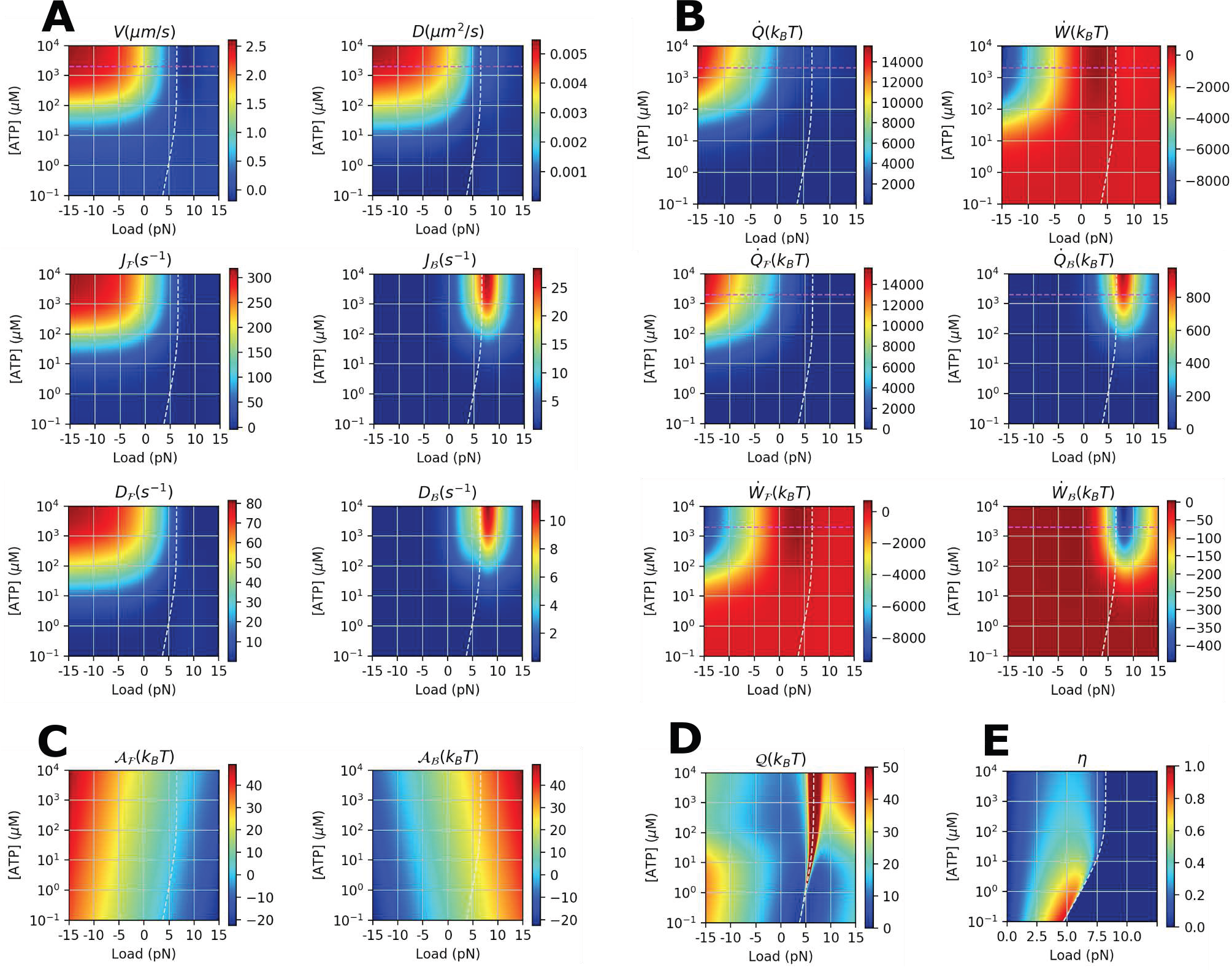
Various physical properties of KIF17 calculated using 6-state network model (Fig. 1A) at varying *f* and [ATP]. **A.** Transport properties (J, D). **B.** Heat 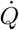 and work production *Ẇ*. **C.** Thermodynamic affinity 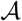, **D.** 𝒬(*f*, [ATP]). **E.** Power efficiency 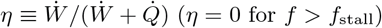. The white dashed lines indicate the stall condition.

**FIG. S8.**
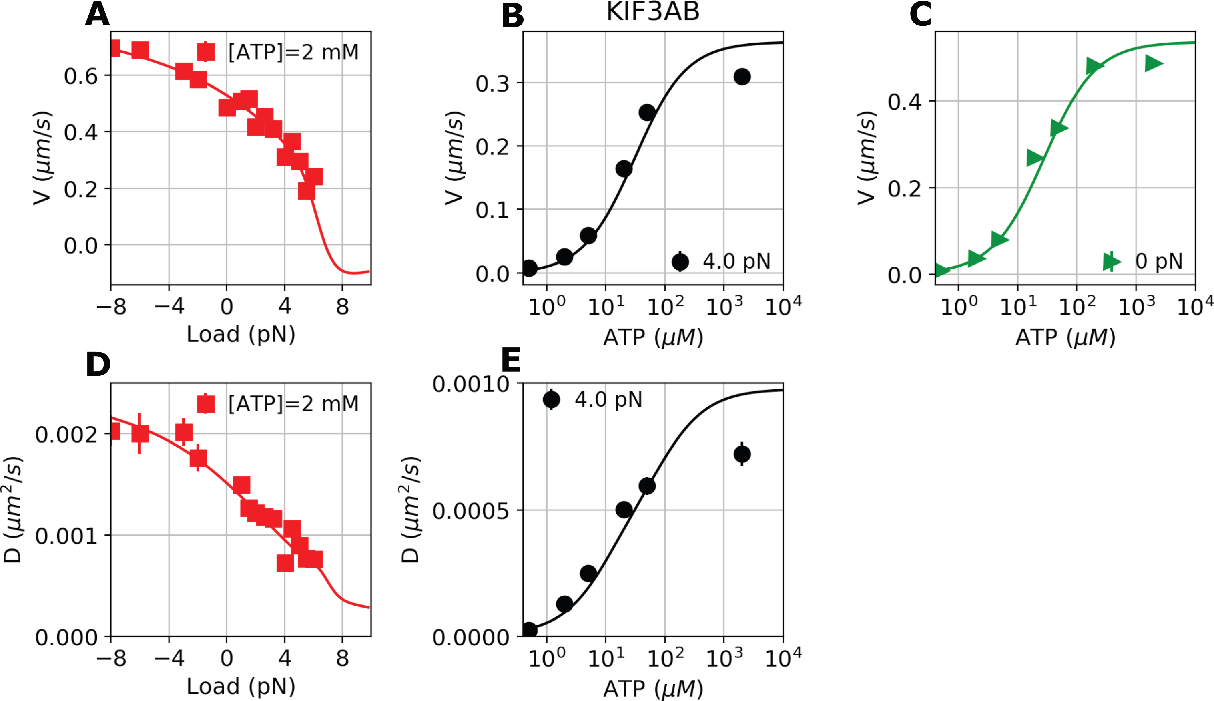
Motility data of heterotrimeric kinesin-2 (KIF3AB) [32] and their theoretical fits (solid lines).

**FIG. S9.**
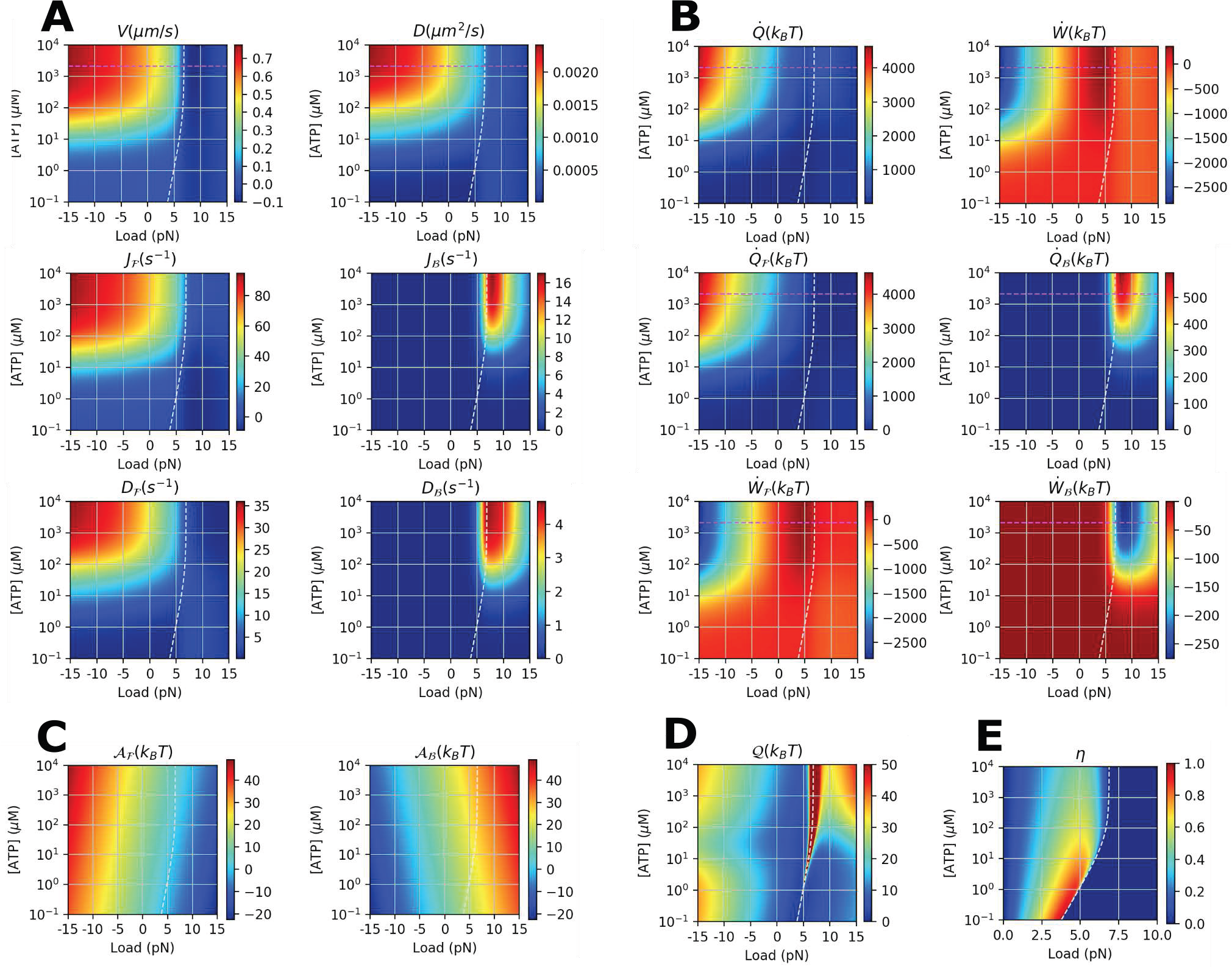
Various physical properties of KIF3AB calculated using the 6-state network model (Fig. 1A) at varying *f* and [ATP]. **A.** Transport properties (*J*, *D*). **B.** Heat 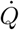 and work production *Ẇ*. **C.** Thermodynamic affinity 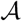. **D.** 𝒬(*f*, [ATP]). **E.** Power efficiency 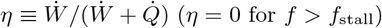. The white dashed lines indicate the stall condition.

**FIG. S10.**
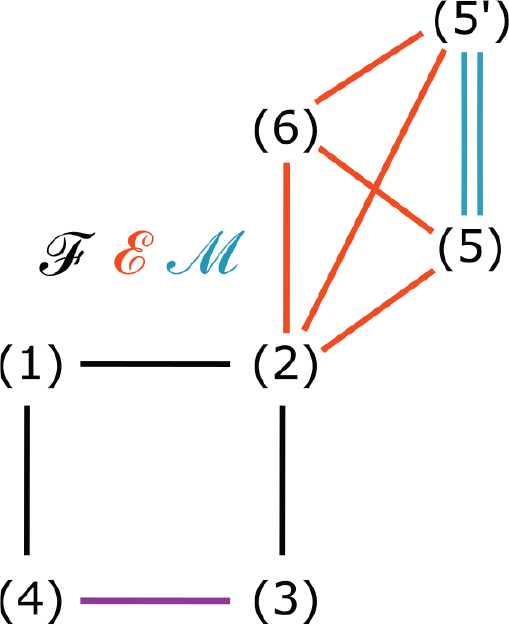
Augmented kinetic network model of myosin-V. Each line represents a reversible kinetics.

**FIG. S11.**
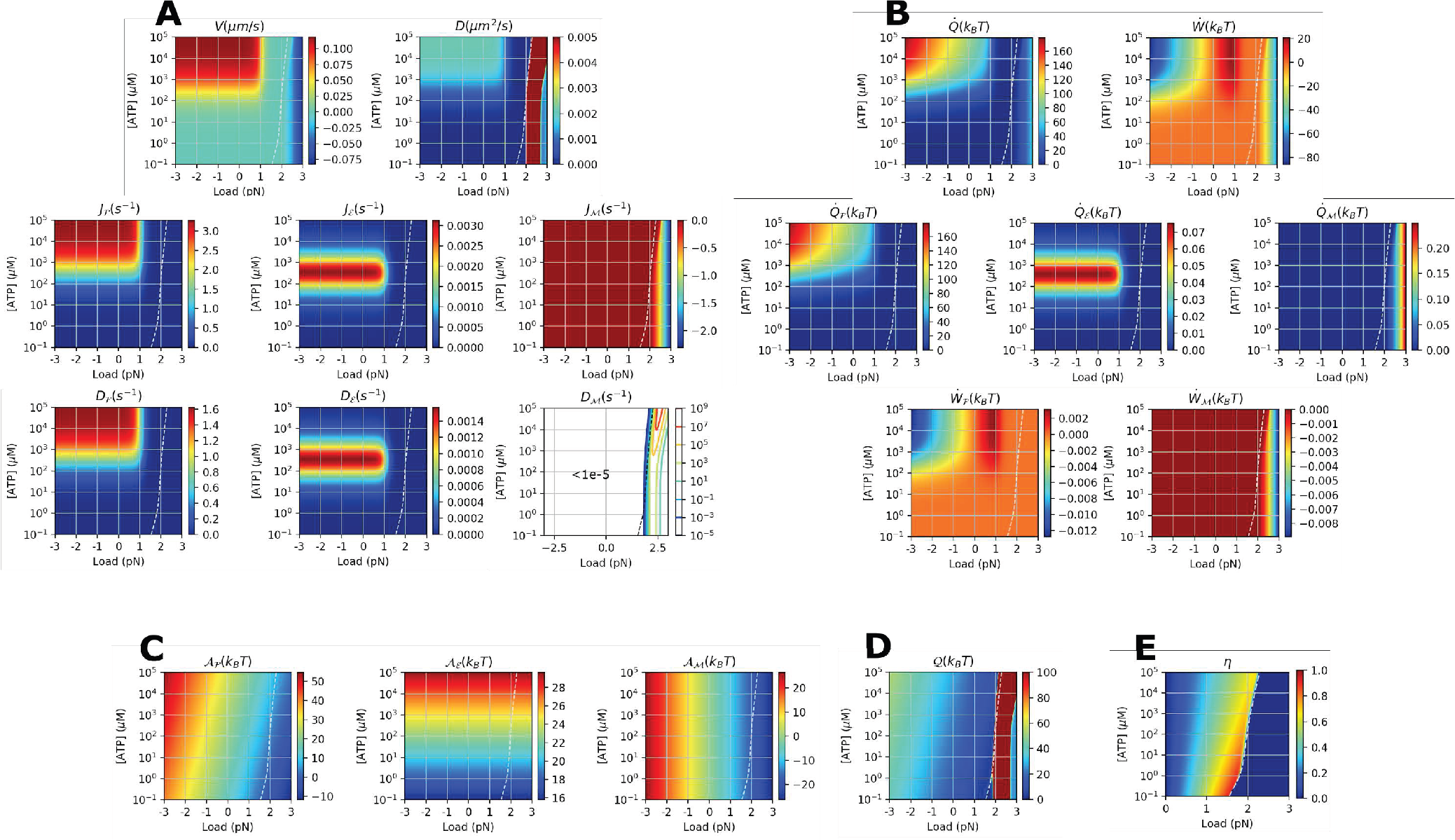
Various physical properties of myosin-V calculated using the multi-cyclic model [38] at varying *f* and [ATP]. [ADP] = 70 *μ*M and [Pi] = 1 mM condition was used for the calculation. **A.** Transport properties (*J*, *D*). **B.** Heat 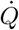 and work production *Ẇ*. **C.** Thermodynamic affinity 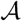. **D.** 𝒬(*f*, [ATP]). **E.** Power efficiency *N* = 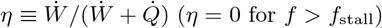. The white dashed lines indicate the stall condition.

**FIG. S12.**
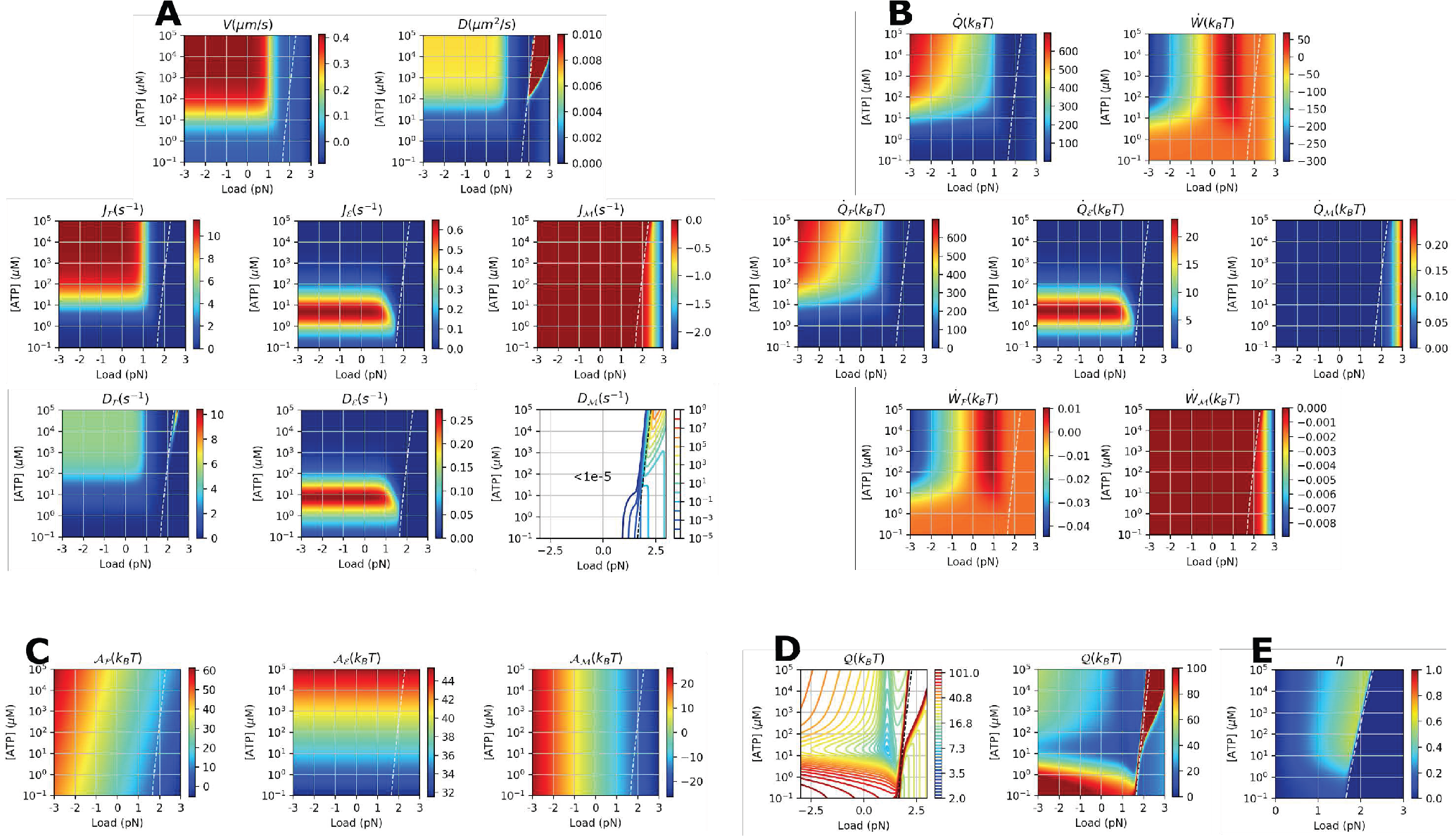
Various physical properties of myosin-V calculated using the multi-cyclic model [38] at varying *f* and [ATP]. [ADP] = 0.1 *μ*M and [P_i_] = 0.1 *μ*M condition was used for the calculation. **A.** Transport properties (*J*, *D*). **B.** Heat 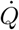 and work production *Ẇ*. **C.** Thermodynamic affinity 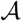. **D.** 𝒬(*f*, [ATP]). **E.** Power efficiency 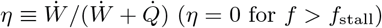. The white dashed lines indicate the stall condition.

**FIG. S13.**
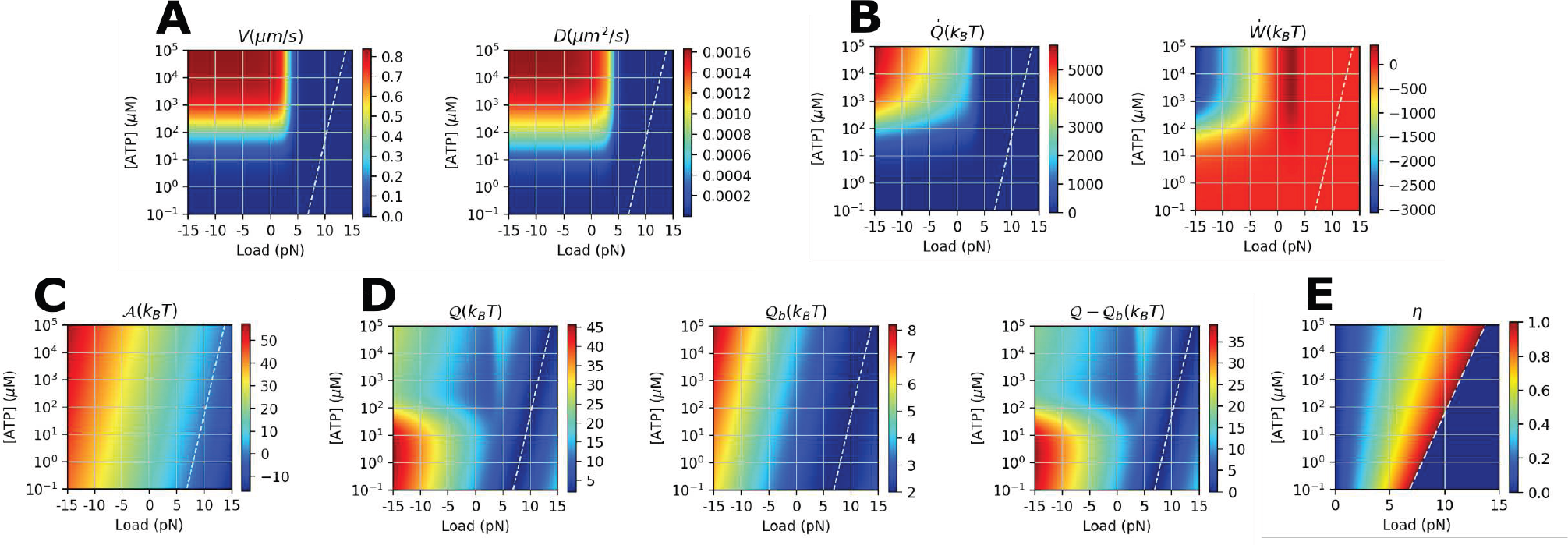
Various physical properties of dynein monomer calculated using (*N* = 9)-unicyclic model [39] at varying *f* and [ATP]. [ADP] = 70 *μ*M and [P_i_] = 1 mM condition was used for the calculation. **A.** Transport properties (*V*, *D*). **B.** Heat 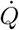 and work production *Ẇ*. **C.** Thermodynamic affinity 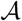. **D.** 𝒬(*f*, [ATP]), 𝒬_*b*_(*f*, [ATP]), and Δ𝒬(*f*, [ATP]). **E.** Power efficiency 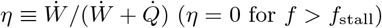. The white dashed lines indicate the stall condition.

**FIG. S14.**
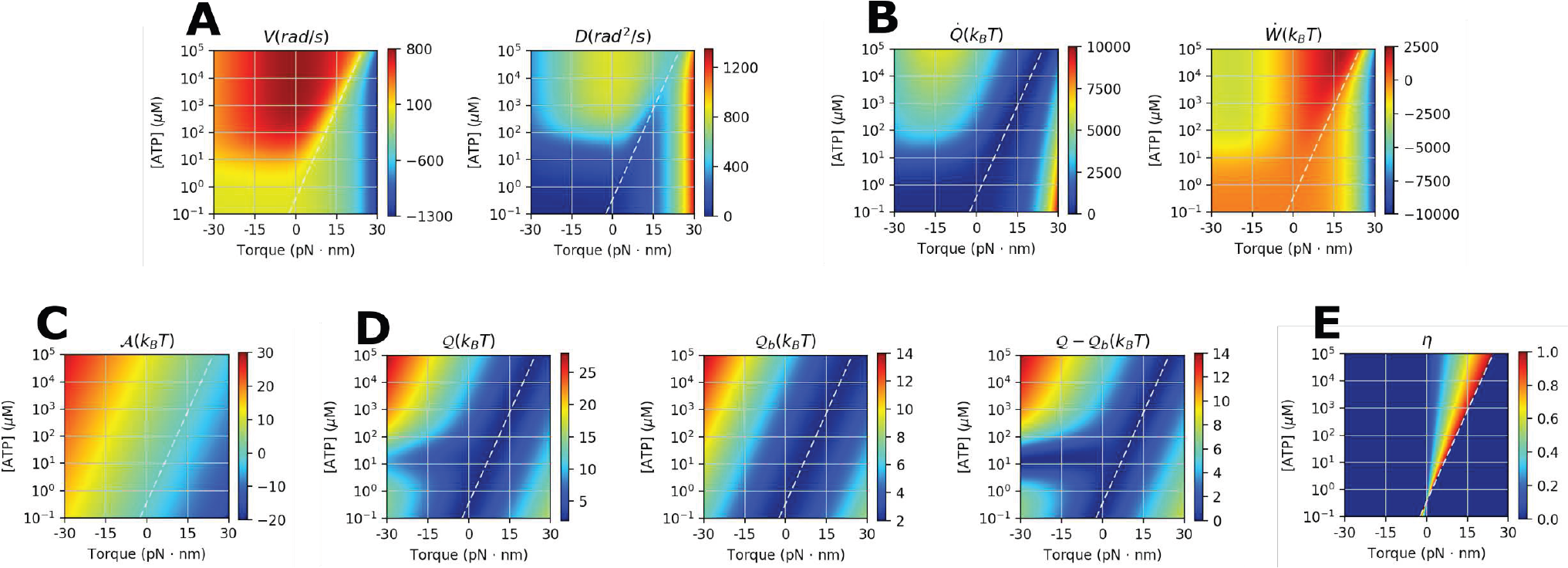
Various physical properties of F_1_-ATPase calculated using (*N* = 2)-unicyclic model (Fig. 4C) at varying *f* and [ATP]. [ADP] = 70 *μ*M and [P_i_] = 1 mM condition was used for the calculation. **A.** Transport properties (*V*, *D*). **B.** Heat 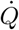 and work production *Ẇ*. **C.** Thermodynamic affinity 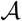. **D.** 𝒬, 𝒬_*b*_ (Eq.S40), and their difference Δ𝒬 = 𝒬−𝒬_*b*_. **E.** Power efficiency 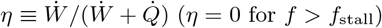. The white dashed lines indicate the stall condition.

**FIG. S15.**
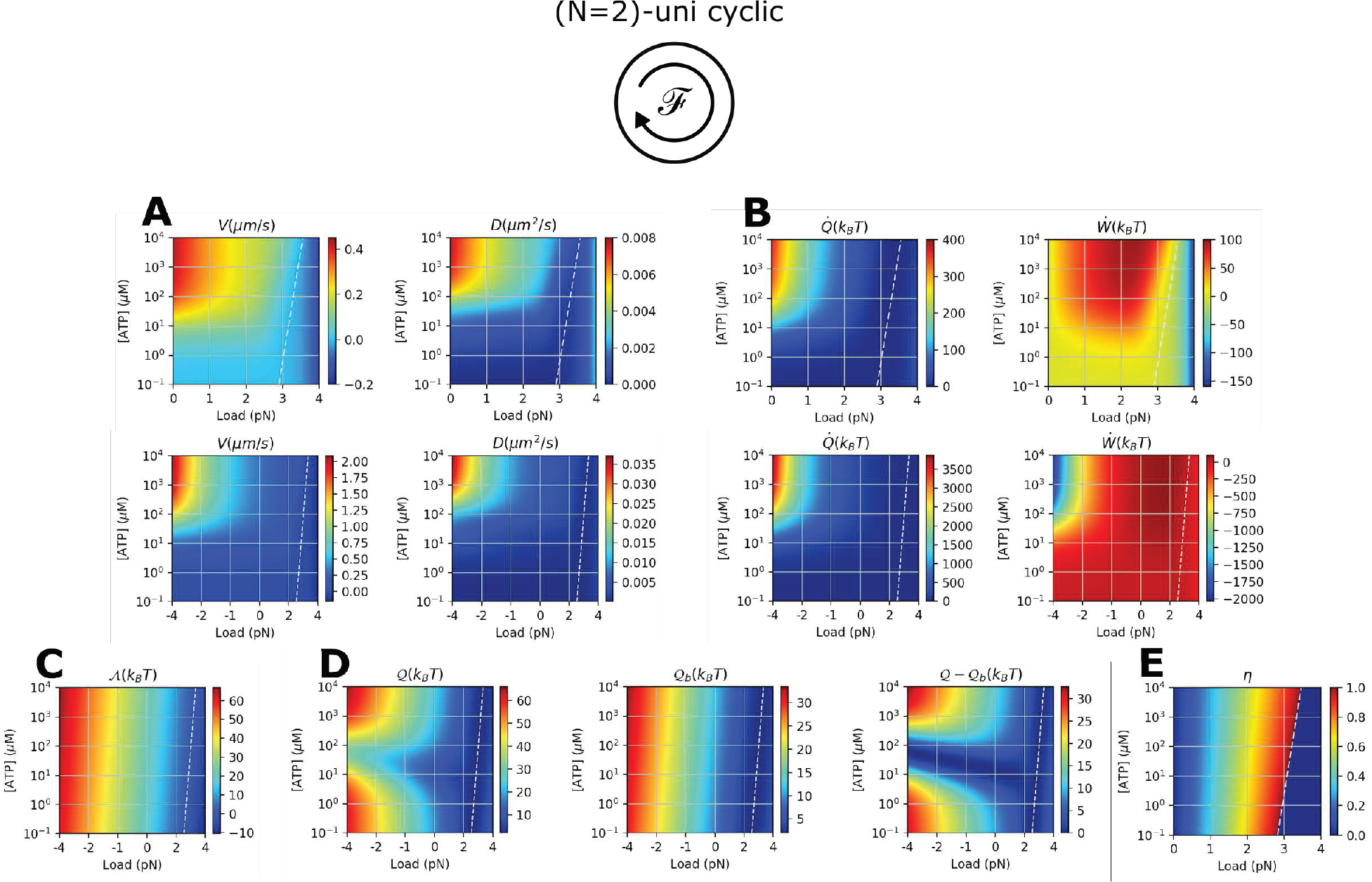
Various physical properties of myosin-V calculated using (*N* = 2)-unicyclic model [41] at varying *f* and [ATP]. **A.** Transport properties (*V*, *D*). Same data are shown twice over the different range of *f* for clarity. **B.** Heat 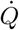 and work production *Ẇ*. Same data are shown twice over the different range of *f* for clarity. **C.** Thermodynamic affinity 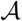. **D.** 𝒬, 𝒬_*b*_ (Eq.S40), and their difference Δ𝒬 = 𝒬−𝒬_*b*_. **E.** Power efficiency 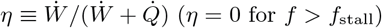. The white dashed lines indicate the stall condition.

